# Efficient and accurate near telomere-to-telomere haplotype reconstruction of diploid genomes

**DOI:** 10.64898/2026.05.20.726711

**Authors:** Yuansheng Liu, Yichen Li, Jialu Xu, Zhongzheng Tan, Wenhai Zhang, Long Wang, Luohao Xu, Jiawei Luo, Xiangxiang Zeng, Alexander Schönhuth, Xiao Luo

## Abstract

Telomere-to-telomere (T2T) and haplotype-resolved assembly are crucial for understanding eukaryotic genomes. For diploid species, this resolution is critical to uncover allelic variations, inheritance patterns, and functional genomic traits. Current scaffolding methods typically employ either sequence-based or graph-based strategies. Sequence-based approaches rely on proximity signals to yield high contiguity, but underutilize assembly graph information, resulting in more structural errors and chromosomal misassignments. Graph-based methods leverage graph topology for higher accuracy but frequently struggle to achieve chromosome-scale contiguity. However, neither strategy alone can overcome its inherent limitations to simultaneously achieve high contiguity and accuracy. To address these challenges, we introduce HapFold, the *first* hybrid scaffolding framework that synergistically leverages the complementary strengths of both graph-based and sequence-based approaches. By integrating the topological accuracy of assembly graphs with the proximity-guided contiguity of sequence models, HapFold achieves highly accurate, chromosome-scale or near-T2T haplotype reconstructions for diploid genomes. Compared to existing methods, HapFold achieves superior assembly quality while accelerating computation by an order of magnitude. Furthermore, in the haplotype reconstruction of diploid genomes using standard Oxford Nanopore Technologies simplex reads, HapFold enables the reconstruction of a greater number of near-T2T assemblies. Our approach provides a robust and scalable solution for the high-fidelity reconstruction of haplotype-resolved diploid genomes.

## Introduction

The completion of the first telomere-to-telomere (T2T) human reference genome [1] has established a new benchmark for genomic completeness, facilitating a transition from fragmented consensus sequences toward the generation of comprehensive, haplotype-resolved diploid assemblies. As most higher animals and plants, including humans, are diploid, a single collapsed genome sequence inherently obscures critical genetic diversity [2], because artificial merging of homologous chromosomes conflates distinct alleles, masks haplotype-specific structural variations, and severs the physical linkages between distant mutations [3]. The missing resolution of the phases severely impedes downstream functional analyses, for example restricting the study of allele-specific expression, cis-regulatory networks, and compound heterozygous mutations in complex disorders [4, 5]. Beyond advancing precision medicine, haplotype-resolved assemblies are essential for decoding the intricate evolutionary histories of animal and plant lineages [6–9]. High-fidelity ploidy aware assembled genomes are also crucial for elucidating the genetic basis of heterosis and accelerating molecular breeding by linking genomic architecture to complex agronomic traits [10–13]. However, accurately separating and reconstructing these highly similar homologous chromosomes across the entire genome remains a fundamental computational challenge in modern genomics [14].

Independent of their advantages in particular applications, haplotype-resolved T2T assemblies establish a canonical standard in computational genomics, because they reflect the optimum both in terms of completeness and evolutionary resolution when reconstructing single genomes. This also explains the great interest in such assemblies as a methodical challenge even before technological advances in genome sequencing raised hopes of fully accomplishing the task. First, encouraged by the advances delivered by third-generation sequencing, seminal methods put emphasis on how to deal with the noise inherent to long reads by employing primary consensus strategies [15–17], which were later followed by haplotype-aware assemblers designed to resolve allelic variations [18, 19]. Subsequently, the highly accurate PacBio High-Fidelity (HiFi) reads catalyzed the development of precision-focused assemblers [20, 21], significantly improving sequence precision and allelic resolution. To completely separate these diploid haplotypes, traditional trio-binning approaches provide highly accurate phasing [11, 22], but their application is decisively constrained by the stringent parental data requirements. Chromatin conformation capture (Hi-C) offers robust haplotype phasing capabilities and has catalyzed the development of various assemblers [23–25]. While Hi-C effectively resolves the phasing challenge, the inherent length limitations of HiFi reads frequently preclude the achievement of T2T-level chromosome reconstruction [26].

To achieve T2T reconstruction, the early algorithms depended on the integration of multiple sequencing modalities. Typically, these methods combined ultra-long Oxford Nanopore Technologies (ONT) reads for extended sequence spanning, and Hi-C reads for haplotype reconstruction. Specifically, advanced assemblers adopted a dual-graph strategy that combines these modalities to assist highly accurate HiFi assemblies in traversing ultra-complex repetitive arrays [27, 28]. However, this approach can compromise the overall assembly quality and incur prohibitive expenses. Recently, error-corrected standard ONT simplex reads have emerged as a more streamlined and efficient paradigm to achieve T2T contiguity [29–31]. However, relying solely on ONT reads cannot achieve haplotype-resolved T2T assembly, requiring external phasing signals from trio-binning or Hi-C.

When primary assemblies fall short of true chromosome-scale contiguity, downstream scaffolding tools are essential to computationally order and orient the fragmented sequences [32, 33]. Leveraging spatial proximity signals from Hi-C [23] or Pore-C [34], these tools offer a highly cost-effective strategy for haplotype resolution. Current scaffolding methodologies utilizing proximity-ligation data are broadly divided into two classes, namely, *sequence-based* and *graph-based approaches* (Table 1). Sequence-based scaffolders initially emerged as existing methodologies designed primarily for haploid or haplotype-collapsed assemblies (e.g., SALSA2 [35], 3D-DNA [36], and YaHS [37]). Subsequently, allele-aware scaffolders (e.g., ALLHiC [38], HapHiC [39]) were developed specifically to disentangle homologous sequences. While these sequence-based tools typically excel at generating assemblies with higher chromosome-level contiguity, they operate independently of the structural information provided by the assembly graph—a network of overlap connections between initial unitigs or contigs generated prior to scaffolding [40]. By processing contigs as discrete linear sequences, they inevitably lose the read-level overlaps and topological connectivity provided in the initial assembly stage [41]. Due to the lack of such information, even allele-aware tools may encounter structural errors, leading to inter-haplotype misjoins and the erroneous fusion of homologous chromosomes. Furthermore, these methods typically require base-level sequence-to-sequence alignments for Hi-C mapping, rendering them computationally expensive and slow.

**Table 1:** Characteristic summary of representative assembly tools. Scaffolding indicates whether the tool includes a dedicated scaffolding module. Haplotype aware denotes the tool’s capability to process and maintain haplotype phasing. T2T level indicates the ability of the scaffolding tool to successfully reconstruct near-T2T scaffolds or contigs. Sequence-based refers to tools that assemble contigs relying solely on sequence-to-sequence Hi-C contact signals. Graph-based indicates that the tool performs local Hi-C signal integration directly upon an assembly graph structure. Application specifies if the tool is designed for specialized tasks, with “General” representing broad, general-purpose usage. Dependency notes whether the tool requires external auxiliary software to complete its pipeline.

| Method | Scaffolding | Haplotype aware | T2T level | Sequence-based | Graph-based | Application | Dependency |
| --- | --- | --- | --- | --- | --- | --- | --- |
| HapFold | Yes | Yes | Yes | Yes | Yes | General | No |
| hifiasm | No | Yes | Yes | No | Yes | General | No |
| pstools | Yes | Yes | No | No | Yes | Cancer | No |
| HapHiC | Yes | Yes | No | Yes | No | General | Yes |
| YaHS | Yes | No | No | Yes | No | General | Yes |
| SALSA2 | Yes | No | No | Yes | No | General | Yes |
| 3D-DNA | Yes | No | No | Yes | No | General | Yes |

In contrast, graph-based approaches alleviate this computational burden by utilizing highly efficient *k*-mer-based sequence-to-graph alignment strategies. However, while these recent methods leverage assembly graph topology to guide phasing—–thereby mitigating localized structural errors—–they often suffer from an inherent algorithmic limitation: they rely on graph topology solely for the phasing step and lack a dedicated graph-based scaffolding mechanism. Consequently, their continuity is strictly bounded by graph connectivity [42]. When topological edges are severed—typically at ultra-complex repeats or centromeres—these graph-dependent frameworks halt [43]. Lacking the ability to utilize broad spatial constraints to span the different components of the disjoint assembly graph, they remain fundamentally constrained in their capacity to achieve true chromosome-scale continuity [44]. For instance, although pstools [45] incorporates graph topology, its haplotype reconstruction framework exhibits clear limitations when resolving complex chromosomal regions mainly because it puts emphasis on cancer genome analysis. Ultimately, graph-based methods face limitations in generating accurate chromosome-scale haplotypes inherent to their mode of operation.

We introduce HapFold ([Hap]lotype-aware Scaf[Fold]er), *the first hybrid scaffolding framework to integrate both graph-based and sequence-based paradigms*. HapFold takes as input unitig graphs generated by state-of-the-art long-read assemblers (e.g., hifiasm [29]) from either PacBio HiFi or ONT simplex reads, alongside long-range chromatin ligation data such as Hi-C or Pore-C, and outputs phased, chromosome-scale to near-T2T assemblies for diploid genomes. Within this seamless synergistic framework, the unitig graph provides high-confidence sequence units and the global topological relationships among them, whereas Hi-C or Pore-C supplies long-range haplotype and spatial information for resolving homologous chromosomes. By jointly using these complementary sources of evidence, HapFold connects, orders, orients, and phases unitigs within a hybrid scaffolding framework, thereby improving both contiguity and haplotype consistency. A decisive contribution of HapFold is that, when applied to assembly graphs generated by hifiasm from standard ONT simplex data, it substantially increases the overall yield of near-T2T haplotypes while preserving phasing accuracy. More remarkably, HapFold also produces a comparable number of near-T2T chromosomes from HiFi-derived unitig graphs, suggesting that this hybrid scaffolding can extend HiFi assemblies toward T2T-scale haplotype reconstruction. Because conventional assembly metrics may overlook long-range scaffold errors and phasing-disruptive structural defects in diploid T2T reconstruction [46–48], we introduce alignment-based metrics to quantify their physical scale and haplotype impact. These metrics allow us to assess not only whether HapFold produces near-T2T chromosomes, but also whether these haplotypes are reliable in both structure and phasing [49–53]. Ultimately, HapFold establishes a robust and efficient new standard for diploid genome scaffolding, facilitating accurate T2T haplotype reconstruction to advance population genomics, clinical genetics, and evolutionary biology.

## Results

### Overview of the HapFold workflow

HapFold introduces a hybrid scaffolding paradigm designed to seamlessly couple the structural precision of graph-based topology with the extensive contiguity of sequence-based methodologies. As illustrated in Fig. 1a, the complete HapFold pipeline encompasses four primary stages. Specifically, read-mapping, graph-refining, and phasing steps are predominantly executed within the graph domain to ensure structural accuracy. Subsequently, the scaffolding step leverages the exact topological boundaries provided by the graph, synergistically combining them with macroscopic sequence-based proximity signals to ultimately assemble chromosome-scale diploid genomes. To ensure clarity, we use ‘contigs’ or ‘unitigs’ to refer to all initial scaffolding inputs, whereas ‘scaffolds’ exclusively denotes the finalized output sequences constructed by ordering, orienting, and bridging these inputs.

**Fig. 1.**
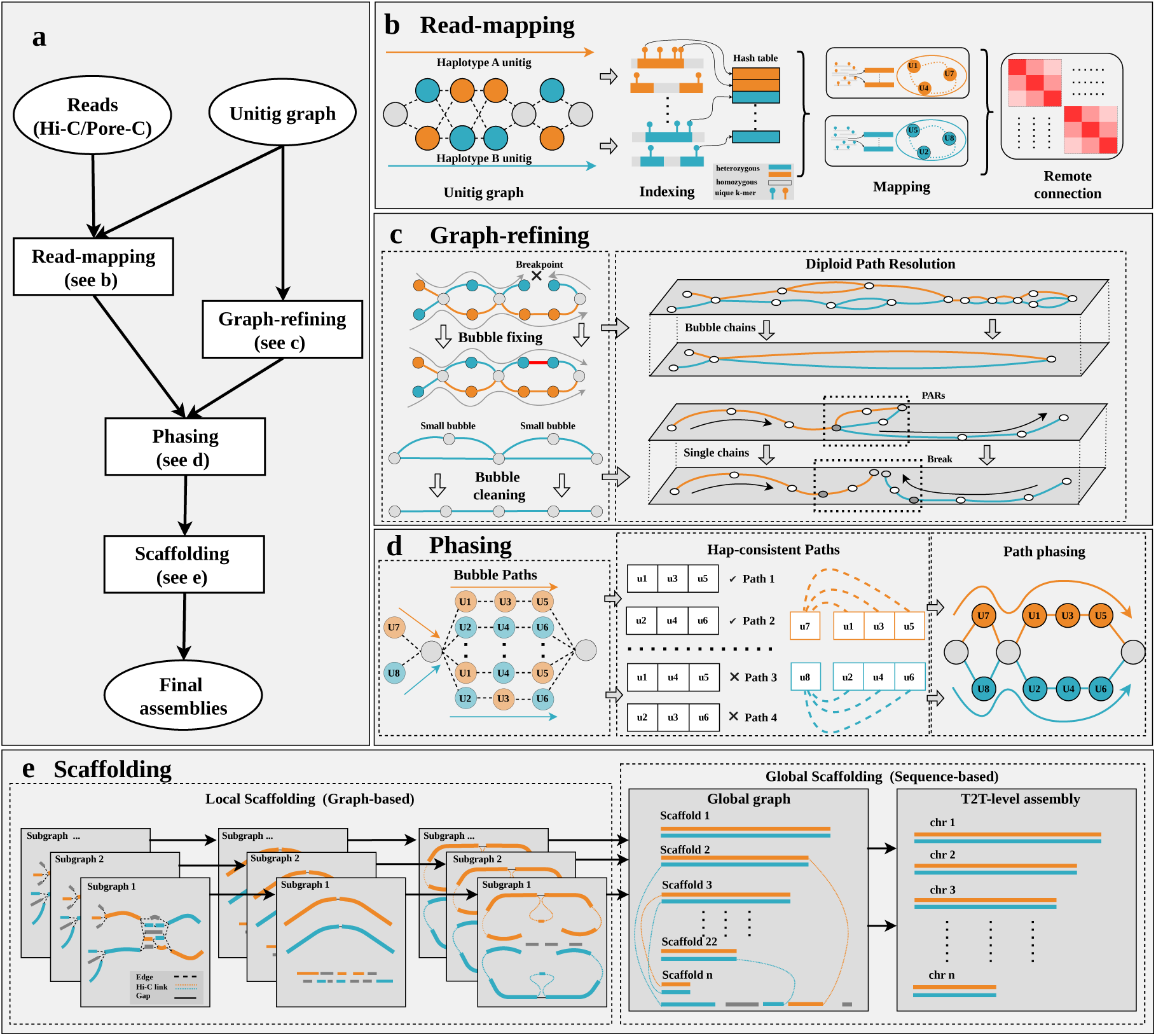
Overview of the HapFold framework for haplotype-resolved diploid assembly. **a**, The integrated workflow of HapFold. Taking a unitig graph (generated by hifiasm or similar assemblers) and proximity-ligation data (Hi-C or Pore-C) as input, the framework executes four core modules: Read-mapping, Graph-refining, Phasing, and Scaffolding. **b**, Haplotype-aware read mapping. Arrows indicate the workflow direction. In the Unitig graph, nodes represent assembled unitig sequences. Orange and cyan indicate distinct haplotypes. Grey denotes homozygous regions. Dashed lines represent initial overlap connections between unitigs. For indexing, unique *k*-mers from heterozygous and homozygous sequences are cataloged in a hash table. During mapping, proximity-ligation read pairs (short fragments linked by dashed lines) align to their corresponding unitigs via the indexed *k*-mer markers. In Remote connection, a signal pairing matrix between unitigs is derived from the proximity-ligation data. **c**, Assembly graph structural refinement. Operating on the initial unitig graph, the bubble fixing stage resolves existing breakpoints, followed by bubble cleaning to consolidate small, fragmented bubbles. In the diploid path resolution stage, dashed lines delineate the graph topology before and after the resolution process. Bubble chains represent the graph architecture generated from homologous regions, while single chains correspond to single-haplotype segments. Additionally, the pseudoautosomal regions (PARs) are specifically identified within the curated graph structure. **d**, Haplotype phasing via proximity signals. Intra-bubble trajectories are resolved into Hap-consistent Paths, where checkmarks denote selected segments and dashed lines represent inter-bubble connectivity. Final haplotype-resolved assemblies are generated during Path phasing via topological extension, following the direction indicated by arrows. **e**, Hierarchical chromosome-scale scaffolding. The scaffolding process is divided into local scaffolding and global scaffolding stages. Within the local scaffolding stage, the sequential panels (from left to right) illustrate the scaffolding of phased contigs within each independent subgraph. During the global scaffolding stage, the global graph represents the integration and further scaffolding of scaffolds merged from all subgraphs. Guided by the target chromosome number (chr *n*), this process ultimately generates the final chromosome-level assembly.

The workflow initiates with two primary inputs: an unphased unitig graph and raw proximityligation reads, such as Hi-C or Pore-C (Fig. 1b). To efficiently process spatial data, HapFold constructs a hash index using unique *k*-mers that appear exactly once within the unitigs, allowing paired proximity-ligation reads to be mapped directly onto the identified sequences. Unique *k*-mers naturally highlights heterozygous loci. Furthermore, a hash-based indexing mechanism constructs these precise signal connections, ensuring the process is highly computationally efficient and entirely dependency-free (Table 1).

To further enhance the overall contiguity of the assembly, HapFold performs systematic graph optimization (Fig. 1c). The initial optimization involves bubble fixing to resolve topological breakpoints that impede phasing, followed by bubble cleaning to remove artifactual small bubbles, thereby yielding a structurally purified graph. Following this refinement, the framework executes diploid path resolution. Rather than applying a uniform simplification, the framework systematically partitions the graph into bubble chains and single chains to accommodate distinct biological scenarios. Bubble chains are utilized to resolve standard homologous chromosomes, where allelic differences manifest as alternative paths within the graph. Conversely, single chains are employed for regions of extreme heterozygosity, such as sex chromosomes, where divergent sequences may fail to form closed bubbles. Notably, these single chains often encapsulate pseudoautosomal regions (PARs, Extended Data Fig. 1). HapFold identifies and separates these PARs based on specific graph structural features, ensuring the correct linkage of the resulting haplotype paths. This structural categorization guarantees that subsequent phasing and scaffolding decisions are grounded in a biologically representative topological substrate.

Building upon this refined graph, HapFold utilizes the spatial information provided by the remote connections to accurately resolve the correct topological linkages both within and between bubbles (Fig. 1d). A common pitfall in haplotype phasing is the interference of signals between homologous chromosomes, which frequently leads to phase switching. HapFold leverages remote connections to iteratively trace intra- and inter-bubble signal linkages to resolve the complete phased paths, a strategy that effectively suppresses noise from homologous regions.

Finally, HapFold executes hierarchical scaffolding (Fig. 1e) to achieve chromosome-scale reconstruction. During the local scaffolding stage, the algorithm first extracts a pair of continuous paths within each subgraph to serve as the backbones. Subsequently, nodes that were bypassed or fragmented during initial path resolution are reattached to their optimal positions on the backbone utilizing Hi-C/Pore-C contact frequencies. This approach is termed graph-based scaffolding because it relies on the reliable structural skeleton of the graph topology and uses proximity-ligation signals to guide the recovery of fragmented nodes. Subsequently, HapFold transitions to a sequence-based global scaffolding module. At this stage, the majority of the generated scaffolds are explicitly paired. Because these scaffolds are constrained by the graph topology, this paired structure effectively prevents the erroneous fusion of homologous chromosomes. Ultimately, HapFold employs a sequence model to order and join these scaffolds based on remote connection signals. By integrating both graph-based and sequence-based methods, HapFold can achieve high structural accuracy and T2T-level contiguity Table 1.

### Datasets

We selected four representative biological datasets (Supplementary Table 1), including one human sample (HG002), two avian samples (Chicken1 and Chicken2 [40]), and one porcine sample (Pig). Specifically, the human HG002 genome was selected because it provides a complete, Q100 diploid human genome benchmark [54]. Achieving near-perfect accuracy across its telomere-to-telomere sequence, it serves as an ideal truth set for exactly quantifying assembly continuity and structural fidelity. The avian samples (Chicken1 and Chicken2) were chosen to test the algorithm’s ability to resolve highly fragmented microchromosomes. The Pig genome [55] was included to represent a complex mammalian genome. Overall, these selected datasets encompass a broad spectrum of structural characteristics, with genome sizes ranging from 1.05 to 3.00 Gb, heterozygosity rates varying from 0.22% to 0.76%, and diploid chromosome numbers (2*n*) spanning from 38 to 78. Our evaluation utilized a diverse combination of sequencing modalities—comprising PacBio HiFi, standard ONT simplex reads, Hi-C, and Pore-C data—demonstrating the framework’s broad applicability across various data types.

### Benchmarking framework for diploid scaffolding

We evaluated HapFold against a comprehensive suite of state-of-the-art tools (Table 1), including established sequence-based Hi-C scaffolders (YaHS [37], HapHiC [39], SALSA2 [35], and 3D-DNA [36]), the graph-based utility pstools [45], and hifiasm [29].

An inherent limitation in evaluating scaffolders arises during the initial assembly stage. As explicitly noted in the hifiasm study [25], the assembler may place paternal and maternal contigs from different chromosomes into a single partition. This represents an innate ambiguity in proximity-ligation phasing (such as Hi-C), while it can successfully partition haplotypes, it cannot inherently assign their parental origin without additional pedigree data. Consequently, relying solely on a single pre-phased result for subsequent downstream scaffolding will inevitably lead to phase switch errors. To overcome this evaluation bias, we adopted the approach utilized by Zeng et al. [39], by merging the haplotype-resolved contigs from hifiasm into a single mixed-sequence dataset. This dataset served as the unified input for all baseline sequence scaffolders, whereas graph-based tools (HapFold and pstools) directly utilized the unphased unitig graph. This mixed-haplotype input paradigm ensures the subsequent construction of more accurate haplotype-resolved assembly results.

Beyond establishing a fair starting baseline, accurate benchmarking also requires a rigorous ground truth to evaluate structural fidelity. Current evaluation methodologies for haplotype-resolved assemblies predominantly rely on alignments against chimeric reference genomes. However, this approach inadvertently masks inter-haplotype mismatches, as aligning to a collapsed haploid reference fails to penalize the incorrect joining of paternal and maternal sequences, thereby falsely inflating the perceived accuracy of the tool. To more accurately evaluate the scaffolding results, we adopted a stringent benchmarking framework based on alignment to a complete, fully phased diploid reference [29]. This evaluation approach more precisely reflects the assembly capability of each scaffolding tool.

### Comprehensive evaluation using exising assembly metrics

HapFold successfully overcomes the historical trade-off between sequence contiguity and structural fidelity in diploid genome scaffolding. Across diverse datasets, it consistently achieves an optimal balance—delivering highly contiguous assemblies while strictly preserving phasing accuracy and structural integrity (Table 2). By seamlessly integrating graph topology with sequence mapping, HapFold fundamentally outperforms both purely sequence-based and purely graph-based competitors.

**Table 2:** Performance comparison of Hi-C-based scaffolding methods. All metrics are reported for the diploid assembly (sum of both haplotypes). Within each dataset block, the internal horizontal line separates graph-based methods (top) from sequence-based methods (bottom). Size: total assembled diploid genome size. # of scaffold (scaffold count): the total number of assembled sequences. NGA50: N50 calculated using reference-aligned blocks, providing a robust measure of structural continuity and accuracy. GF (genome fraction): percentage of the reference genome successfully covered by the assembly. DR (duplication ratio): ratio of total aligned bases in the assembly to the total covered bases in the reference. # of MS (misassembled scaffolds): number of scaffolds containing structural assembly errors (relocations, inversions, or translocations). Genome fraction, NGA50, duplication ratio, and number of misassembled scaffolds were comprehensively evaluated using the tool QUAST [50]. HE (hamming error): proportion of incorrectly phased alleles within phase blocks. SE (switch error): rate of phase switches between adjacent heterozygous loci. Both phasing metrics were evaluated using the *k*-mer-based tool yak [22]. ‘-’ indicates that the tool failed to generate results for this dataset.

| Dataset | Assembler | Size<br>(Gb) | GF<br>(%) | # of<br>Scaffold | NGA50<br>(Mb) | DR | # of<br>MS | HE<br>(%) | SE<br>(%) |
| --- | --- | --- | --- | --- | --- | --- | --- | --- | --- |
| HG002<br>HiFi+Hi-C | HapFold | 5.95 | 98.93 | 104 | 99.12 | 1.00 | 52 | 3.77 | 0.92 |
|  | hifiasm | 5.99 | 99.46 | 631 | 74.20 | 1.00 | 75 | 1.54 | 0.97 |
|  | pstools | 8.97 | 88.23 | 826 | 49.46 | 1.68 | 94 | 1.35 | 0.91 |
|  | HapHiC | 5.99 | 99.52 | 426 | 91.86 | 1.00 | 67 | 6.88 | 0.97 |
|  | YaHS | 5.99 | 99.50 | 269 | 92.24 | 1.00 | 69 | 14.90 | 0.97 |
|  | SALSA2 | 5.99 | 99.48 | 540 | 87.01 | 1.00 | 101 | 3.94 | 0.97 |
|  | 3D-DNA | - | - | - | - | - | - | - | - |
| Chicken1<br>HiFi+Pore-C | HapFold | 2.21 | 97.82 | 273 | 29.02 | 1.05 | 50 | 0.84 | 0.26 |
|  | hifiasm | 2.15 | 98.76 | 1043 | 20.26 | 1.01 | 53 | 0.77 | 0.54 |
|  | pstools | 2.35 | 78.00 | 334 | 14.49 | 1.37 | 67 | 15.65 | 0.41 |
|  | HapHiC | 2.15 | 97.70 | 940 | 21.49 | 1.02 | 67 | 2.09 | 0.54 |
|  | YaHS | 2.15 | 95.29 | 490 | 31.19 | 1.04 | 188 | 23.35 | 0.54 |
|  | SALSA2 | 2.15 | 98.90 | 777 | 20.26 | 1.01 | 135 | 10.42 | 0.54 |
|  | 3D-DNA | 2.15 | 98.81 | 3586 | 5.55 | 1.01 | 53 | 0.67 | 0.54 |
| Chicken2<br>HiFi+Hi-C | HapFold | 2.16 | 93.50 | 628 | 18.75 | 1.08 | 101 | 2.67 | 1.20 |
|  | hifiasm | 2.12 | 97.99 | 787 | 14.35 | 1.01 | 52 | 1.85 | 1.27 |
|  | pstools | 2.15 | 70.06 | 288 | 13.29 | 1.41 | 88 | 3.56 | 1.22 |
|  | HapHiC | 2.12 | 93.45 | 229 | 19.03 | 1.06 | 62 | 7.89 | 1.28 |
|  | YaHS | 2.12 | 92.09 | 329 | 19.06 | 1.06 | 126 | 8.39 | 1.29 |
|  | SALSA2 | 2.12 | 98.00 | 582 | 14.35 | 1.01 | 114 | 4.58 | 1.28 |
|  | 3D-DNA | 2.12 | 97.19 | 3603 | 3.41 | 1.03 | 53 | 1.47 | 1.24 |
| Pig<br>HiFi+Hi-C | HapFold | 5.39 | 95.37 | 335 | 83.91 | 1.11 | 75 | 5.30 | 2.38 |
|  | hifiasm | 5.14 | 97.21 | 721 | 34.35 | 1.04 | 112 | 2.66 | 2.51 |
|  | pstools | 5.46 | 82.90 | 372 | 31.48 | 1.25 | 210 | 3.03 | 2.52 |
|  | HapHiC | 5.14 | 95.98 | 307 | 65.12 | 1.05 | 74 | 6.30 | 2.52 |
|  | YaHS | 5.14 | 97.59 | 164 | 83.91 | 1.04 | 66 | 18.39 | 2.52 |
|  | SALSA2 | 5.14 | 97.20 | 489 | 34.35 | 1.04 | 145 | 9.06 | 2.52 |
|  | 3D-DNA | 5.14 | 98.31 | 991 | 24.16 | 1.03 | 111 | 2.65 | 2.51 |
| HG002<br>ONT+Hi-C | HapFold | 6.00 | 99.19 | 56 | 133.57 | 1.00 | 33 | 3.89 | 1.00 |
|  | hifiasm | 6.00 | 99.72 | 209 | 105.77 | 1.00 | 45 | 1.68 | 0.96 |
|  | pstools | 18.70 | 80.36 | 1316 | 54.51 | 3.86 | 229 | 4.06 | 2.11 |
|  | HapHiC | 6.00 | 99.73 | 156 | 105.77 | 1.00 | 44 | 4.55 | 0.96 |
|  | YaHS | 5.99 | 99.65 | 118 | 105.77 | 1.00 | 47 | 4.29 | 0.97 |
|  | SALSA2 | 5.99 | 99.73 | 186 | 105.77 | 1.01 | 49 | 7.22 | 0.97 |
|  | 3D-DNA | 5.99 | 99.71 | 211 | 105.77 | 1.01 | 46 | 1.68 | 0.96 |

#### Assembly continuity and genomic completeness

To evaluate assembly continuity and structural accuracy, we primarily employed the NGA50 metric, which accounts for structural misassemblies by breaking scaffolds at alignment breakpoints, alongside the total scaffold count, the number of misassembled scaffolds (which explicitly quantifies erroneously joined sequences), and the duplication ratio. Across the evaluated datasets, HapFold delivered highly competitive NGA50 metrics while substantially reducing total scaffold counts. For instance, HapFold improved NGA50 by 31% to 144% relative to hifiasm and consolidated the assemblies by reducing the total scaffold count by 20% to 82%. Crucially, HapFold accomplished this extensive contiguity while effectively controlling the number of misassembled scaffolds, avoiding the severe structural degradation observed in other highly contiguous outputs. This exceptional balance is remarkably evident in the HG002 ONT dataset. While all competing scaffolders stalled at the baseline NGA50 of 105.77 Mb, HapFold uniquely propelled the NGA50 to an unprecedented 133.57 Mb, simultaneously achieving the lowest scaffold count (56) and minimizing misassemblies (33).

In contrast, while sequence-based methods often achieve higher contiguity, graph-based methods typically maintain better structural accuracy. However, neither category matches the comprehensive performance balance of HapFold. Among sequence-based tools, legacy methods like SALSA2 and 3D-DNA exhibited fragmented outputs, failing to meaningfully improve NGA50 over the hifiasm contigs. Notably, 3D-DNA generated over 3,500 disconnected scaffolds in the highly heterozygous Chicken datasets. More recent sequence-based tools demonstrated better contiguity but suffered from structural flaws. For example, although YaHS achieved an NGA50 comparable to HapFold on the Chicken2 dataset, it lacks adequate diploid phasing capability. This limitation led to a 3.45% to 5.90% drop in genome fraction compared to hifiasm, and concurrently generated the highest number of misassembled scaffolds. HapHiC, utilizing a haplotype-aware algorithm, achieved good contiguity with fewer severe structural errors than YaHS. Nevertheless, its overall performance still fell short of HapFold. Conversely, while the graph-based tool pstools natively leverages structural topology, it struggles with global contiguity. Because it is optimized for targeted genomic regions rather than whole-genome scaffolding, it produced significantly lower genome fractions, fragmented NGA50 scores, and anomalous duplication ratios.

Notably, a substantial disparity was observed between the N50 (Supplementary Table 2) and NGA50 metrics across various scaffolding tools, which stems from the presence of numerous potential structural errors within the assembly results. N50 reflects mere physical length, whereas NGA50 represents the contiguous length strictly aligned to a reference without errors. Consequently, relying solely on N50 often inflates perceived continuity and obscures misassemblies. This limitation is particularly evident in sequence-based scaffolders, which frequently exhibit a pronounced gap between N50 and NGA50 (a low NGA50/N50 ratio). This discrepancy indicates that sequence-based methods are highly susceptible to introducing structural errors, primarily because they lack the topological constraints necessary to accurately differentiate between homologous alleles. For example, in the HG002 dataset, YaHS generated an NGA50/N50 ratio of merely 57.3% (92.24 Mb / 160.91 Mb), while HapHiC produced a restricted ratio of 64.0%. In stark contrast, HapFold maintained a robust NGA50/N50 ratio of 70.8% (99.12 Mb / 139.97 Mb), proving its ability to extend contiguity without compromising structural fidelity.

However, the NGA50 metric also has inherent limitations, as it represents continuity strictly at a single, median-centric percentile threshold. To evaluate structural connectivity comprehensively across the entire assembly scale, we generated NGAx distribution curves (Fig. 2). The NGAx trajectories clearly demonstrate that HapFold outperforms competing tools across the vast majority of assembly ranges. For instance, HapFold exhibits pronounced superiority from the 20th to the 60th percentiles in HG002, and dramatically outperforms all other tools from the 0 to the 50th percentile in the Pig dataset. For the Chicken datasets, the absence of a complete T2T reference genome naturally compresses the NGAx trajectories. While YaHS appears to achieve superficially higher NGAx values in these avian datasets, this elevation does not reflect true continuity. Rather, it exposes the tool’s inadequate diploid phasing capability, which easily concatenates multiple homologous sequences together.

**Fig. 2.**
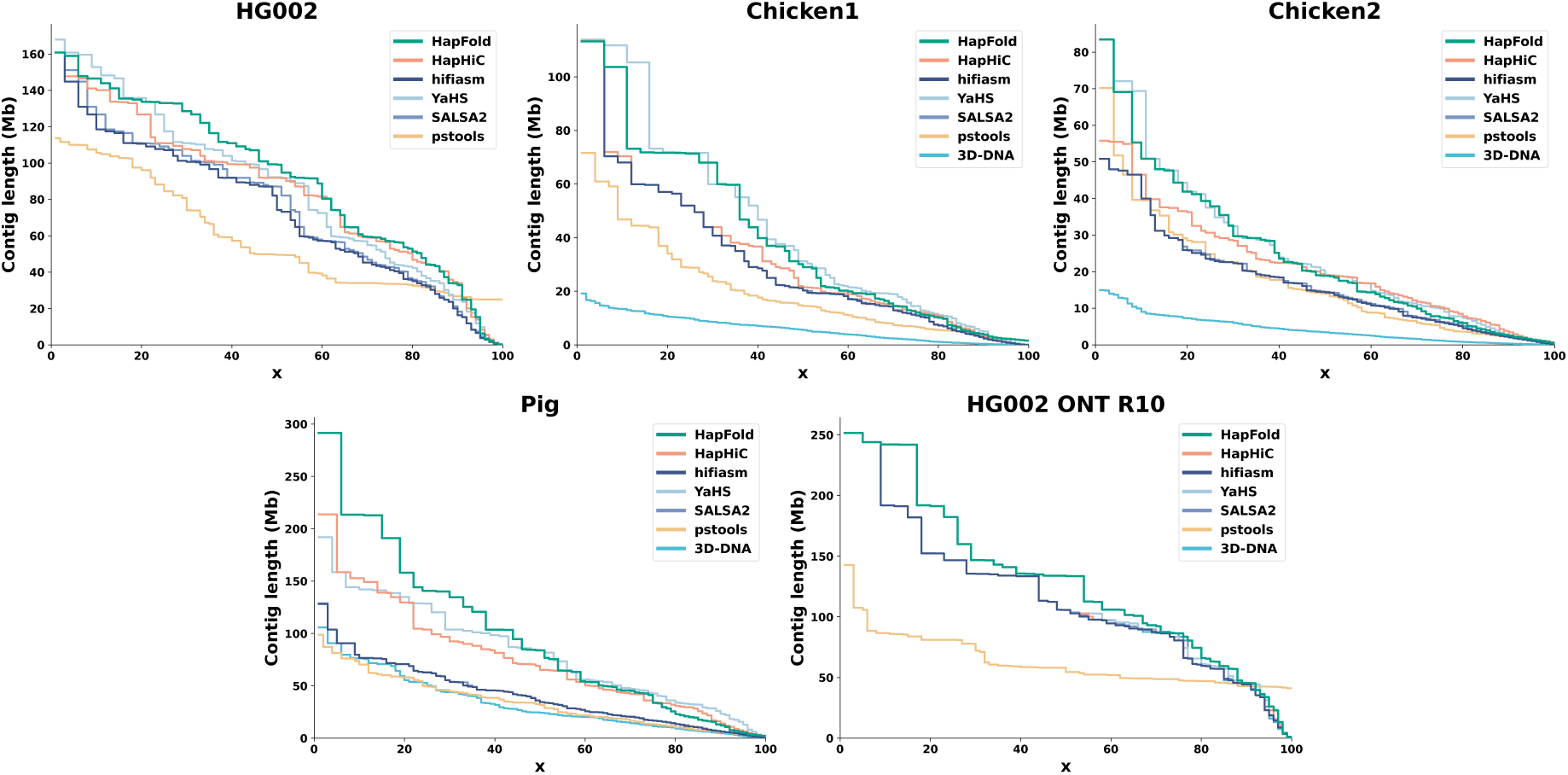
NGA*x* connectivity curves across five benchmark datasets. Each subplot displays the aligned contig length (Mbp) on the y-axis against the cumulative percentage of the genome (*x*) on the x-axis. The results encompass five datasets: HG002, Chicken 1, Chicken 2, Pig, and HG002-ONT Simplex Reads. Curves for six scaffolding methods (HapFold, hifiasm(Hi-C), HapHiC, YaHS, SALSA2, pstools and 3D-DNA) are color-coded as indicated in the legends. A higher curve and a more gradual horizontal decay represent greater effective assembly continuity, where the NGA*x* values account for structural accuracy by considering only correctly aligned genomic segments. Discontinuities or rapid drops in the curves reflect points where the cumulative assembly length is composed of increasingly smaller or misassembled scaffolds.

YaHS effectively trades higher assembly errors for inflated continuity metrics. Excluding this over-scaffolding, the NGAx curves confirm that HapFold consistently delivers the most structurally sound and optimal true continuity across all evaluated datasets.

#### Haplotype phasing accuracy

To evaluate haplotype phasing accuracy, we measured Hamming and switch error rates across assemblies. Hamming error quantifies the ability to correctly separate maternal and paternal sequences, while switch error measures the phase consistency within continuous blocks. Overall, HapFold demonstrated superior phasing accuracy, consistently maintaining significantly lower Hamming error rates compared to all sequence-based scaffolders. For example, in the HG002 dataset, HapFold reduced the Hamming error to 3.77%, objectively outperforming HapHiC (6.88%) and YaHS (14.90%). Although the graph-based tool pstools recorded a lower Hamming error, its framework is specifically tailored for targeted cancer genome analysis; consequently, it produces severely fragmented assemblies and is unsuitable for comprehensive chromosome-scale scaffolding.

The lower Hamming error achieved by HapFold stems from its hybrid architecture. By utilizing graph topological constraints, HapFold strictly prevents the erroneous fusion of homologous sequences. In contrast, sequence-based scaffolders directly process linear contigs, which leads to varying degrees of increased phasing errors. Notably, HapFold also produces a lower switch error because its graph-based mechanism actively corrects internal phase errors within contigs. For example, in the Chicken1 dataset, HapFold reduced the switch error from 0.54% with hifiasm to 0.26%. Conversely, sequence-based scaffolders merely reorder and orient existing contigs; as a result, they typically maintain the initial baseline switch error and are unable to improve intra-contig phasing.

Furthermore, this phasing accuracy is corroborated by the discrepancy between NGA50 and N50 metrics. Sequence-based methods frequently exhibit a large NGA50/N50 gap alongside elevated Hamming errors. This correlation confirms that sequence-based methods suffer from higher haplotype phasing error rates, as their assembly contiguity often incorporates a higher frequency of structural errors. Pore-C technology captures long-range, multi-way chromatin contacts, providing broader spatial constraints than standard pairwise Hi-C. When processing Pore-C data (the Chicken1 dataset), HapFold demonstrated distinct advantages. It delivered the highest structurally sound contiguity while keeping both Hamming and switch errors lower than competing scaffolders. The success of HapFold on Pore-C data is primarily attributed to its *k*-mer-based mapping algorithm. Compared to traditional alignment methods, this *k*-mer strategy is highly efficient at mapping longer, multi-contact Pore-C reads directly to the unitig graph.

To directly visualize the phasing purity of individual scaffolds, we generated yak-based scatter plots representing parental-specific *k*-mer counts (Supplementary Fig. 2). In these plots, high-fidelity scaffolds align tightly with the horizontal and vertical axes. HapFold scaffolds cluster more closely along the axes, whereas other tools show more dispersion toward the diagonal, indicating a higher degree of inter-homolog mixing.

Importantly, these scatter plots highlight distinct genomic signatures, particularly for sex chromosomes. Allosomes extend significantly further along the axes because they possess substantially more unique *k*-mers than autosomes. This visual evidence reveals an inherent limitation of the aggregate Hamming error metric. Because the metric simply sums all misassigned *k*-mers, the overall Hamming error is disproportionately influenced by the massive weight of sex chromosomes.

### Structural fidelity of scaffolded genomes

Relying solely on Hamming error or *k*-mer distributions to evaluate scaffold quality presents certain limitations. Because *k*-mer-based metrics primarily classify sequences into maternal or paternal origins, they exhibit limited sensitivity toward inter-chromosomal misjoins, particularly when such structural errors occur between segments from the same parental haplotype. Furthermore, the true error rates within autosomes are often disproportionately masked by sex chromosomes, which possess a higher abundance of lineage-specific *k*-mers. Similarly, while widely used metrics like NGA50 effectively measure contiguity by penalizing non-contiguous blocks, they fail to capture internal discrepancies regarding sequence ordering and orientation. Moreover, standard discrete counts of “misassembled scaffolds” only record the frequency of error events (i.e., breakpoints), failing to quantify the actual genomic length compromised by these scaffolding misjoins. Therefore, for assemblies with available high-quality references, direct global sequence alignment remains a much more accurate and comprehensive approach to evaluating structural fidelity [1, 2, 54].

#### Characterizing structural misassemblies in scaffolds

To comprehensively capture these alignment-based discrepancies, it is crucial to recognize that structural errors in scaffolds pose significantly more severe risks than those in contig assemblies. While contig misassemblies typically arise from localized sequence overlaps, scaffolding errors entail the erroneous concatenation of massive, pre-assembled contigs. Because these contigs represent substantial genomic blocks, ordering and orientation misjoins induce extensive disruptions in macro-collinearity. Consequently, a simple tally of discrete error events severely underestimates the total impact of scaffolding failures. To address this, we directly align the scaffolds to the reference genome and calculate the total number of bases in misassembled scaffolds [50]. This approach enables us to capture the extent of chromosomal disruption caused by erroneous contig linkages.

By systematically aligning the benchmark assemblies against their respective references, we identified a substantial number of large-scale structural misassemblies specifically introduced during the scaffolding process. To comprehensively characterize these scaffolding-induced anomalies, we explicitly categorize them into three primary structural errors and two chromosome-level errors. The structural errors comprise Inversion errors (Extended Data Fig. 2a), which occur when a scaffolder assigns an incorrect relative orientation to a contig; Duplication errors (Extended Data Fig. 2b), where identical regions from homologous chromosomes are incorrectly collapsed into a single tandem array; and Relocation errors (Extended Data Fig. 2c). A Relocation error is defined by alignment discrepancies where the starting coordinate of a subsequent segment maps to an earlier position than the termination of the preceding segment—indicating improper repositioning—or when contiguous sequences are erroneously separated by artificial gaps. Furthermore, we define two severe chromosome-level misassemblies: Chromosome phase errors (Extended Data Fig. 2d), characterized by erroneous crossover switches between maternal and paternal homologous sequences within a single scaffold, and inter-chromosomal Translocation errors (Extended Data Fig. 2e), which involve the aberrant fusion of segments from entirely different, non-homologous chromosomes. Importantly, these structural misassemblies often manifest concurrently. For instance, Duplication errors are frequently accompanied by Chromosome phase errors. This co-occurrence typically arises when a scaffolder erroneously stitches together homologous segments from both the maternal and paternal alleles at the same genomic locus—an artifact that is particularly prevalent among sequence-based tools.

As demonstrated in our prior metric evaluations, HapHiC stands out as the most competitive haplotype-aware scaffolder among existing methods, exhibiting the fewest overall errors. Consequently, we focus our high-resolution structural analysis on a direct comparison between HapFold and HapHiC. To visualize these complex structural fidelity issues, we aligned the HG002 scaffolds generated by both tools against the high-quality Q100 HG002 reference genome (Fig. 3a). This alignment plot directly illustrates how each segment of the assembled scaffolds aligns to the maternal (red) and paternal (blue) reference sequences.

**Fig. 3.**
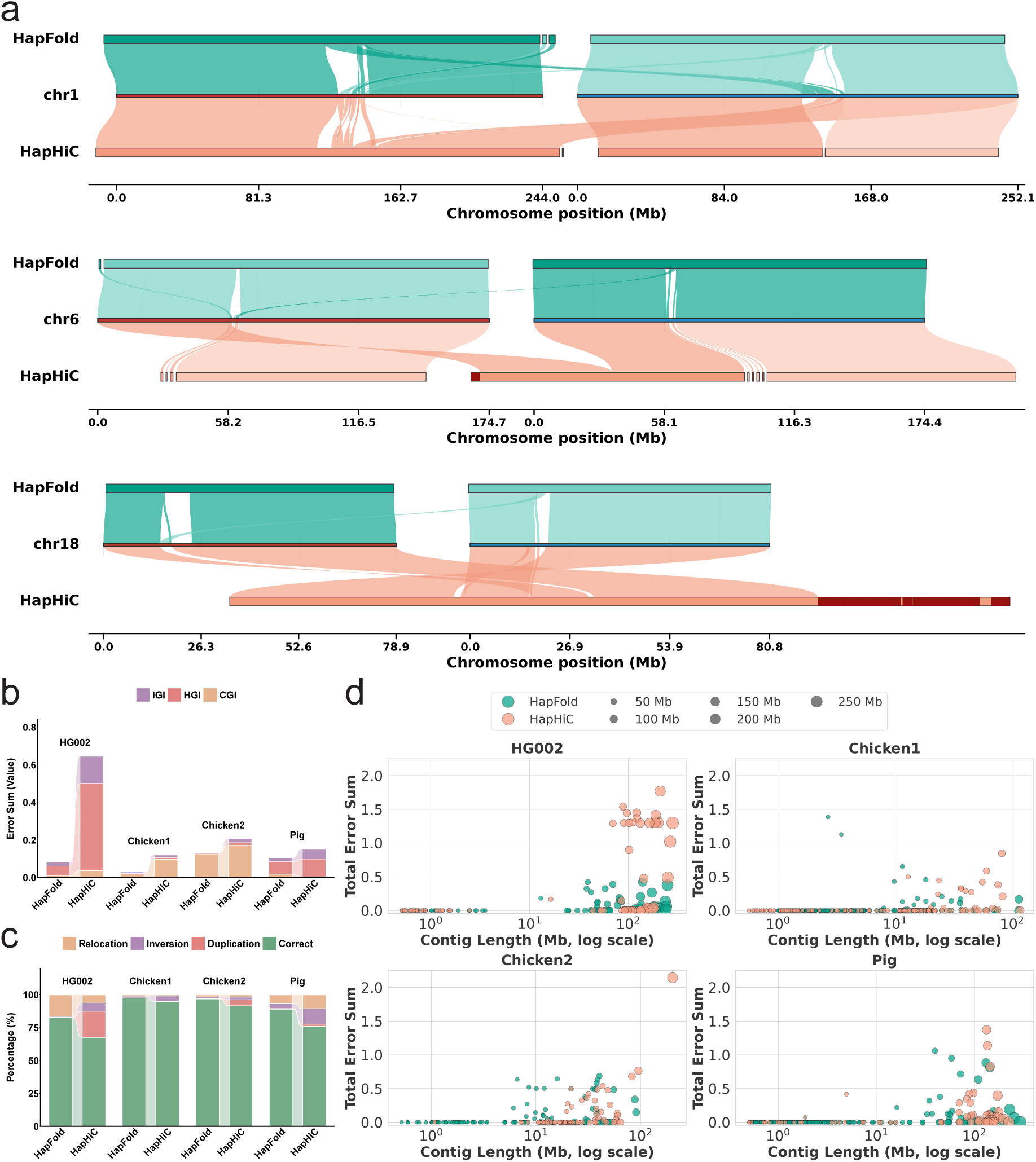
Structural error analysis and phasing accuracy benchmarks across diverse datasets. **a**, Comparative alignment visualization for chromosomes 1, 6, and 18. Red segments highlight regions misaligned to non-target chromosomal areas (translocations). Maternal and paternal haplotypes are displayed on the left and right sides of each coordinate axis, respectively. **b**, Proportion of structural error types. The bar charts represent the percentage of total sequence length classified as correct or attributed to specific structural errors, including relocation, inversion, and duplication. **c**, Cumulative error metrics. The error sum reflects the integrated values of the Intra-chromosomal Gini impurity (IGI), Haplotype Gini impurity (HGI), and Chromosome Gini impurity (CGI), representing structural, phase, and translocation errors, respectively. Each metric is calculated as a proportion of the sequence length, with a maximum theoretical sum of 3.0. **d**, Distribution of individual sequence errors. Scatter plots display the total error sum for individual scaffolds relative to their sequence length on a logarithmic scale, highlighting the error profile across different species and datasets.

This high-resolution mapping reveals that HapFold exhibits fewer errors compared to HapHiC. On chromosome 1, HapFold achieves a largely complete reconstruction, notwithstanding a minor chromosome switch error in a small middle segment. In contrast, HapHiC displays a more extensive chromosome phase error near the 130 Mb coordinate, where a segment of approximately 120 Mb switches from the maternal to the paternal track. Notably, two distinct scaffold segments align to this same paternal region simultaneously. This occurs because the scaffold incorporates an excessive amount of paternal-specific sequences during assembly, which biases the regional alignment toward the paternal reference. On chromosome 6, HapFold successfully reconstructs both paternal and maternal chromosomes, except for minor gaps in the middle. Conversely, HapHiC erroneously merges maternal and paternal sequences between 0 and 58 Mb. This leads to severe structural inversion and duplication errors intertwined with chromosome phase errors.

On chromosome 18, HapFold demonstrates high structural and haplotypic fidelity. It successfully reconstructs both the maternal and paternal alleles of the chromosome; aside from expected gaps, its alignments remain strictly consistent with the reference tracks. HapFold accurately preserves the correct physical order while significantly minimizing the occurrence of phase crossovers and structural misjoins. In contrast, HapHiC exhibits an increased number of scaffolding errors on this same chromosome. It collapses both haplotypes into a fragmented single scaffold containing various structural errors. Moreover, the dark red blocks at the distal end of this HapHiC scaffold denote the erroneous inclusion of non-homologous chromosomal sequences, resulting in an inter-chromosomal Translocation error. Ultimately, collectively, these examples highlight HapFold’s enhanced capability to independently resolve homologous chromosomes and prevent the complex, large-scale misassemblies often encountered by other tools.

These visual comparisons highlight the inherent limitations of existing sequence-based scaffolding methods. Even advanced allele-aware tools designed for haplotype phasing, such as HapHiC, still exhibit numerous structural misassemblies. This occurs because sequence-based approaches typically process contigs as discrete linear sequences, thereby losing the crucial read-level overlap information established during the initial assembly. Consequently, relying solely on Hi-C contact frequencies is often insufficient to guarantee accurate positional anchoring and relative orientation. In contrast, HapFold addresses these challenges through its unique hybrid scaffolding architecture. By integrating the strict topological constraints of graph-based methods with the long-range extension capabilities of sequence-based approaches, HapFold successfully balances the two: it achieves the low structural error rates characteristic of graph methods while delivering the high chromosome-scale contiguity typical of sequence models.

While scaffolding produces sequences significantly longer than initial contigs, our alignment results demonstrate that a single orientation or ordering mistake during this process can trigger severe, chromosome-scale structural disruptions. Unfortunately, the true magnitude of such topological collapses is easily masked by existing evaluation methods. These tools typically penalize these massive disruptions as merely a single discrete misassembly event, fundamentally ignoring the immense genomic length affected by the error. Given the expansive scale of scaffolded sequences, evaluating scaffolding tools based on the physical length of structural errors—rather than simple event counts—is critical. Because current discrete counting metrics fail to fully capture the true impact of these large-scale topological disruptions, we next introduce several newly quantified metrics. These indices are specifically designed to rigorously measure the exact genomic length compromised by these distinct structural errors.

#### Alignment-based quantification of structural errors

To overcome the aforementioned limitations of *k*-mer based phasing evaluations and discrete counting metrics, we introduce a comprehensive suite of alignment-based Gini impurity indices [56]—namely, the intra-chromosomal Gini impurity (IGI), the Haplotype Gini impurity (HGI), and the Chromosome Gini impurity (CGI)—to rigorously assess assembly consistency based directly on physical alignment lengths (Extended Data Fig. 2). First, IGI evaluates three distinct types of intra-chromosomal assembly errors: relocations, inversions, and duplications (Extended Data Fig. 2a–c). These structural inconsistencies occur when different segments of a scaffold erroneously align to disparate positions or orientations within the same reference chromosome. Second, rather than relying on sequence signatures, HGI quantifies the severity of haplotype switching (Extended Data Fig. 2d) by calculating the exact genomic length of scaffold regions that alternate between maternal and paternal reference alignments. Finally, CGI measures inter-chromosomal misjoins (Extended Data Fig. 2e), capturing the proportion of errors where a single assembled scaffold incorrectly aligns across multiple distinct chromosomes.As illustrated in Fig. 3b, the stacked bar charts plot the sum of these alignment-based error proportions for the assembly, revealing the true extent of structural misassemblies and haplotype mixing. For each individual metric, the quantified error value is normalized to a maximum of 1. Overall, the structural error evaluation of the assembly results demonstrates that HapFold consistently achieved the lowest error rates across all datasets. Conversely, HapFold exhibited exceptional structural accuracy, generating substantially lower total error sums across the evaluated datasets. Specifically, within the HG002 dataset, HapFold’s total error sum was restricted to approximately one-eighth that of HapHiC, underscoring its robust capacity to prevent structural misjoins. A similar performance advantage was observed in the Pig dataset, where HapFold yielded a highly accurate assembly, whereas HapHiC produced an error sum nearly 1.5 times greater. It should be noted that for the Chicken1 and Chicken2 datasets, due to the absence of available T2T references, we initially generated their assemblies using trio data. These assemblies were subsequently aligned to a closely related reference genome [57] to establish chromosomal assignments. Consequently, inherent inaccuracies or misassemblies within these proxy references may introduce potential baseline biases into the evaluation.

To comprehensively quantify the proportion of structural misassemblies, Fig. 3c illustrates the fraction of total assembly length affected by three distinct structural error types. Beyond phasing consistency, HapFold demonstrates superior macroscopic structural fidelity by maintaining a significantly higher proportion of correctly aligned scaffolds. Specifically, this correctly aligned fraction exceeds 80% in HG002 and approaches 95% in both Chicken1 and Chicken2. Conversely, HapHiC introduces substantial structural misassemblies, most notably characterized by massive duplication errors within the HG002 dataset.

To further investigate the error distributions within individual assembled scaffolds, Fig. 3d presents a scatter plot of the error sums for each scaffold generated by HapFold and HapHiC. In this visualization, both the x-axis position and the node size represent the scaffold length. For longer scaffolds, the error proportion per scaffold in HapFold is markedly lower than that in HapHiC. This is visually evident as HapFold’s scaffolds predominantly cluster in the bottom-right quadrant of the plot, whereas HapHiC’s scaffolds frequently localize to the top-right quadrant, with error sums regularly exceeding 1.0 in datasets such as HG002 and Pig. This spatial distribution clearly indicates that HapHiC suffers from a notably higher error frequency within its extended scaffolds. In stark contrast, HapFold successfully generates comparably massive, chromosome-scale scaffolds while strictly suppressing the associated error sums near the baseline.

#### Genome-wide visualization of structural fidelity

Following the quantitative assessment, we performed a macroscopic visual comparison of the haplotype-resolved assemblies for the HG002 dataset. This assessment utilized the HG002 Q100 reference as an absolute ground truth for physical mapping. Direct alignment of the scaffold assemblies against this reference provides an intuitive evaluation of overall quality and structural fidelity (Fig. 4).

**Fig. 4.**
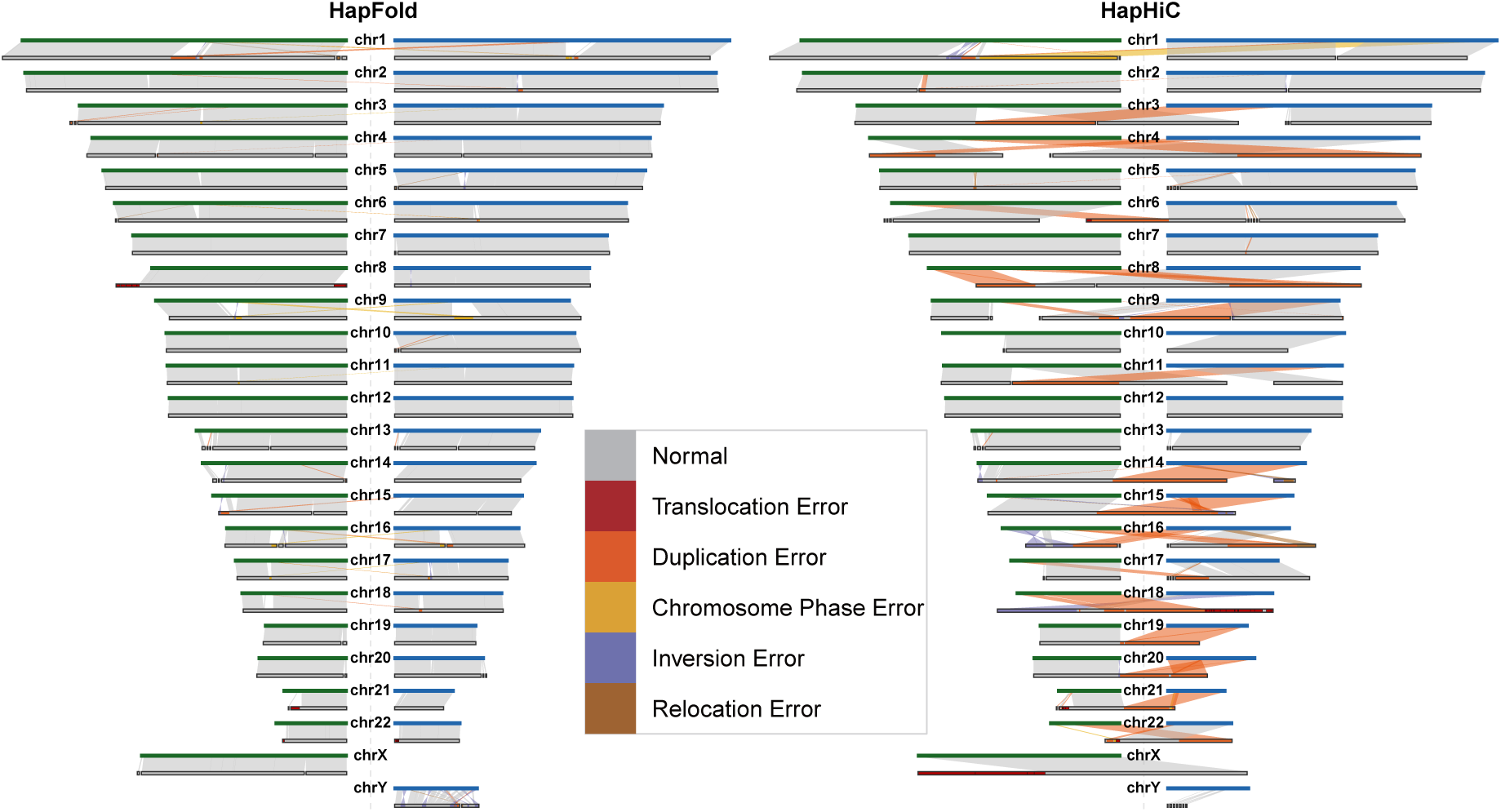
Genome-wide structural alignment of HapFold and HapHiC assemblies using the HG002 T2T Q100 reference. Reference maternal and paternal chromosomes are indicated in green and blue, respectively, and are positioned on the left and right sides of each horizontal track. The assembly scaffolds are represented in grey. Correct collinear alignments are shown as grey ribbons, whereas structural misassemblies are categorized as cross-chromosome translocations (red), inter-homolog duplications (orange), phase switches (yellow), inversions (purple), and intra-chromosome translocations (brown).

As visualized in the genome-wide alignments, HapFold maintains the correct physical sequence order across almost all chromosomes and exhibits virtually no structural misassemblies, particularly avoiding chromosome phase errors. This allows it to generate near-chromosome-scale scaffolds with higher structural fidelity. This is achieved because HapFold utilizes a hybrid strategy that synergistically integrates graph-based and sequence-based scaffolding methods. Specifically, during the initial graph-based processing, HapFold leverages unitig graph topologies—utilizing heterozygous bubble structures that naturally separate homologous alleles—to reconstruct highly symmetric homologous chromosome pairs. Building upon this, the graph-based scaffolding module robustly extends these contigs into longer, well-resolved scaffolds while strictly preventing inter-homolog crossovers. Finally, the sequence-based module utilizes proximity-ligation data to further link the scaffolds that could not be connected within the graph, driving the assembly to a contiguous, chromosome-level resolution while ensuring the symmetry of homologous chromosomes. In contrast, tools like HapHiC rely exclusively on sequence-based scaffolding, making them highly susceptible to the inherent limitations of proximity-ligation data. Because diploid homologous chromosomes occupy nearly identical three-dimensional spatial territories, their Hi-C interaction profiles are often indistinguishable. As a result, HapHiC exhibits varying degrees of structural errors on multiple chromosomes, such as chr1, chr3, chr4, chr8, and chr9. This demonstrates that while sequence-based methods such as HapHiC can construct extended scaffolds, they frequently stitch the physically corresponding maternal and paternal regions into a single collapsed sequence. This inherent methodological limitation inevitably introduces numerous structural misassemblies, primarily manifesting as phase switches and inter-homolog duplications (represented by the yellow and orange ribbons in Fig. 4).

Nevertheless, the alignments reveal specific localized anomalies. Terminal sequence loss or minor misassemblies occur near the telomeres of acrocentric chromosomes, such as chr15, chr21, and chr22. This issue stems from the high biological complexity of ribosomal DNA (rDNA) arrays. These arrays contain ultra-long tandem repeats that often collapse into complex topological tangles during initial HiFi assembly (Extended Data Fig. 1). HapFold utilizes specialized segmentation algorithms to isolate these tangles. However, fragmented unitigs at the boundaries may occasionally mis-scaffold to other acrocentric chromosomes due to their high sequence similarity. The alignments also contain various unaligned gaps on the reference tracks. These gaps do not necessarily indicate missing sequences in the assembly. During competitive alignment against a diploid reference, a scaffold segment may map preferentially to the alternate haplotype. This creates an artificial alignment void on the current haplotype track. Furthermore, other unaligned segments may reside on separate, smaller scaffolds. Some gaps also represent unsequenced regions produced by spatial scaffolding algorithms when bridging disjointed unitigs.

The genome-wide alignment demonstrates high collinearity between the HapFold assembly and the diploid HG002 T2T reference. Notably, the scaffolds generated by HapFold achieve near-complete reconstruction of individual chromosomes, effectively establishing a one-to-one correspondence between a single scaffold and a full chromosome. However, due to the inherent length limitations of HiFi data in spanning highly complex genomic regions, the resulting assemblies inevitably contain varying degrees of gaps. Nevertheless, as evidenced by the comparison with HapHiC, HapFold still achieves better performance in the haplotype-resolved reconstruction of diploid chromosomes, consistently ensuring fundamental structural correctness.

### Robustness evaluation across varying Hi-C sequencing depths

We evaluated the robustness of HapFold across varying Hi-C sequencing depths to assess its sensitivity to proximity-ligation data volume. Current sequence-based scaffolders typically require high Hi-C coverage to achieve accurate ordering and orientation. To characterize this impact, we conducted a gradient benchmarking experiment on the HG002 dataset using four distinct Hi-C depths: 28x, 54x, 76x, and 100x. These results are summarized in Supplementary Table 3; notably, the evaluation at the standard 76x coverage has been previously detailed in Table 2.

As illustrated in Extended Data Fig. 3, we systematically compared the performance of all tools across the different Hi-C depths, focusing on four key metrics: Scaffold Number, NGA50 (Mb), NGA90 (Mb), and Hamming Error (%). First, HapFold consistently produced the lowest Scaffold Number across all tested depths. Compared to the contig outputs of hifiasm, HapFold reduced the total number of scaffolds by approximately 4-to 5-fold on average. This stable reduction indicates that HapFold effectively minimizes assembly fragmentation, reflecting a higher degree of structural completeness regardless of the available Hi-C data volume.

Next, we evaluated NGA50 and NGA90, which measure assembly contiguity at the 50th and 90th percentiles of the genome size, respectively. While NGA50 captures overall macroscopic continuity, NGA90 specifically reflects the connectivity of the more challenging, fragmented tail-end regions of the assembly. HapFold exhibited the highest contiguity in both metrics across all depths. This demonstrates that HapFold successfully integrates a larger number of contigs into extended scaffolds, improving both the overall assembly continuity and the contiguity of smaller, more fragmented sequences. The combination of a minimized scaffold count and enhanced contiguity serves as a strong indicator of HapFold’s structural accuracy.

Finally, we assessed the Hamming Error, which quantifies the phasing accuracy of the assemblies. HapFold maintained a Hamming Error that was only marginally higher than those of the primary contig outputs from hifiasm and the purely graph-based tool, pstools. In contrast, sequence-based methods such as HapHiC demonstrated a stronger dependency on sequencing depth, with their phasing errors exhibiting a gradual decrease only as Hi-C coverage increased. The ability of HapFold to sustain a consistently low Hamming Error across all depth gradients highlights the robust nature of its hybrid approach, proving its capacity to tolerate lower-depth Hi-C data while maintaining high phasing accuracy. Overall, the hybrid methodology of HapFold achieves a phasing accuracy comparable to that of graph-based methods, while simultaneously delivering higher contiguity than existing sequence-based scaffolders. Crucially, it maintains these distinct advantages stably across varying Hi-C sequencing depths.

### Performance across varying heterozygosity in plants

Diploid plant genomes display highly variable heterozygosity—the level of sequence divergence between homologous alleles. Due to the current lack of comprehensive, ground-truth phased data for real plant genomes spanning a broad and continuous heterozygosity spectrum, we utilized simulated plant datasets. We selected multiple diverse, real rice reference genomes and paired them to construct pseudo-diploid genomes. Due to the natural divergence limits among the available high-quality rice reference genomes,the maximum heterozygosity achievable through these pairwise combinations was capped at 1.07%. Based on these pseudo-diploid references, we utilized pbsim2 to simulate HiFi reads and sim3C to generate the corresponding Hi-C spatial data, creating a continuous testing gradient with heterozygosity levels ranging from 0.37% to 1.07%. We benchmarked HapFold against hifiasm and HapHiC, evaluating the performance across the gradient using total scaffold number, NGA50, and NGA90 (Extended Data Fig. 4).

According to our experimental results, initially, as heterozygosity increases from the baseline, we observe an early trend of improvement in assembly contiguity metrics across all assembly tools. This phenomenon suggests that a moderate increase in heterozygosity introduces sufficient sequence divergence, which effectively aids the upstream assembler in distinguishing homologous alleles and forming clear, resolvable bubble structures within the assembly graph. However, this situation indicates that as heterozygosity continues to rise beyond the 0.64% threshold, the higher heterozygosity may introduce complex, highly bifurcated graph topologies and fragmented allelic regions. This increases the processing difficulty for assembly tools, making it more challenging to generate highly contiguous assembly results.

Amidst these complex dynamics, HapFold demonstrates highly stable scaffolding resolution. Across all heterozygosity levels, HapFold consistently maintains the lowest total scaffold count, validating its superior capability to accurately consolidate fragmented segments into continuous architectures. Specifically, HapFold maintains a scaffold count closely aligning with the expected diploid chromosome number of rice (2n = 24), fluctuating narrowly between 23 and 26. In contrast, hifiasm ranges from 30 to 48 contigs, while HapHiC yields 18 to 35 scaffolds overall. This indicates that HapHiC’s scaffold count fluctuates substantially, largely driven by the variations in the initial hifiasm contigs across different heterozygosity levels.

Furthermore, the NGA50 metric reveals that hifiasm and the other evaluated tools have essentially achieved optimal contiguity at the 50th percentile. Consequently, the primary challenge—and the key differentiator of scaffolding performance—lies in the more difficult 90th percentile (NGA90). The NGA90 metric reflects the assembly’s structural continuity across the more fragmented and difficult-to-assemble tail ends of the genome. In this context, HapHiC and hifiasm display nearly identical NGA90 values, indicating that HapHiC fails to enhance contiguity in these challenging regions. Conversely, HapFold demonstrates a clear advantage, improving the NGA90 metric by approximately 20% relative to both HapHiC and hifiasm. Ultimately, these simulation experiments indicate that HapFold delivers stable and accurate chromosome-scale scaffolding across highly variable heterozygosity landscapes, effectively navigating both the phasing advantages of sequence divergence and the structural challenges of extreme allelic complexity.

### T2T-level assembly

Driven by recent algorithmic advancements in primary assemblers such as hifiasm [29, 30], achieving T2T genome assemblies has become increasingly viable. To rigorously evaluate HapFold’s capacity for T2T-level assembly, we conducted comprehensive comparative assessments using both HiFi and standard ONT simplex reads, with each dataset paired with Hi-C data. The resulting scaffolds were then evaluated by aligning them to the HG002 Q100 reference genome [54]. To facilitate downstream scaffolding, telomere-preserved results from hifiasm were utilized as the primary input for both HapFold and HapHiC.

HapFold not only yielded a greater number of T2T-level reconstructions using the ONT dataset, but it also demonstrated exceptional performance on HiFi data (Fig. 5). Furthermore, HapFold consistently achieved a higher number of highly complete chromosomes (defined as T2T-level or achieving ≥ 99% chromosomal completeness) across both sequencing technologies. Following the evaluation criteria established by the Human Pangenome Reference Consortium (HPRC) [2], we define T2T-level assembly as the successful structural recovery of both telomeres for a given chromosome. Since a T2T status emphasizes the reconstruction of telomeres at both ends, it does not necessarily represent the complete sequence spanning the entire chromosomal length. To distinguish actual sequence integrity from the T2T status, we define chromosomal completeness as the genome fraction of individual scaffolds relative to their corresponding reference chromosomes.

**Fig. 5.**
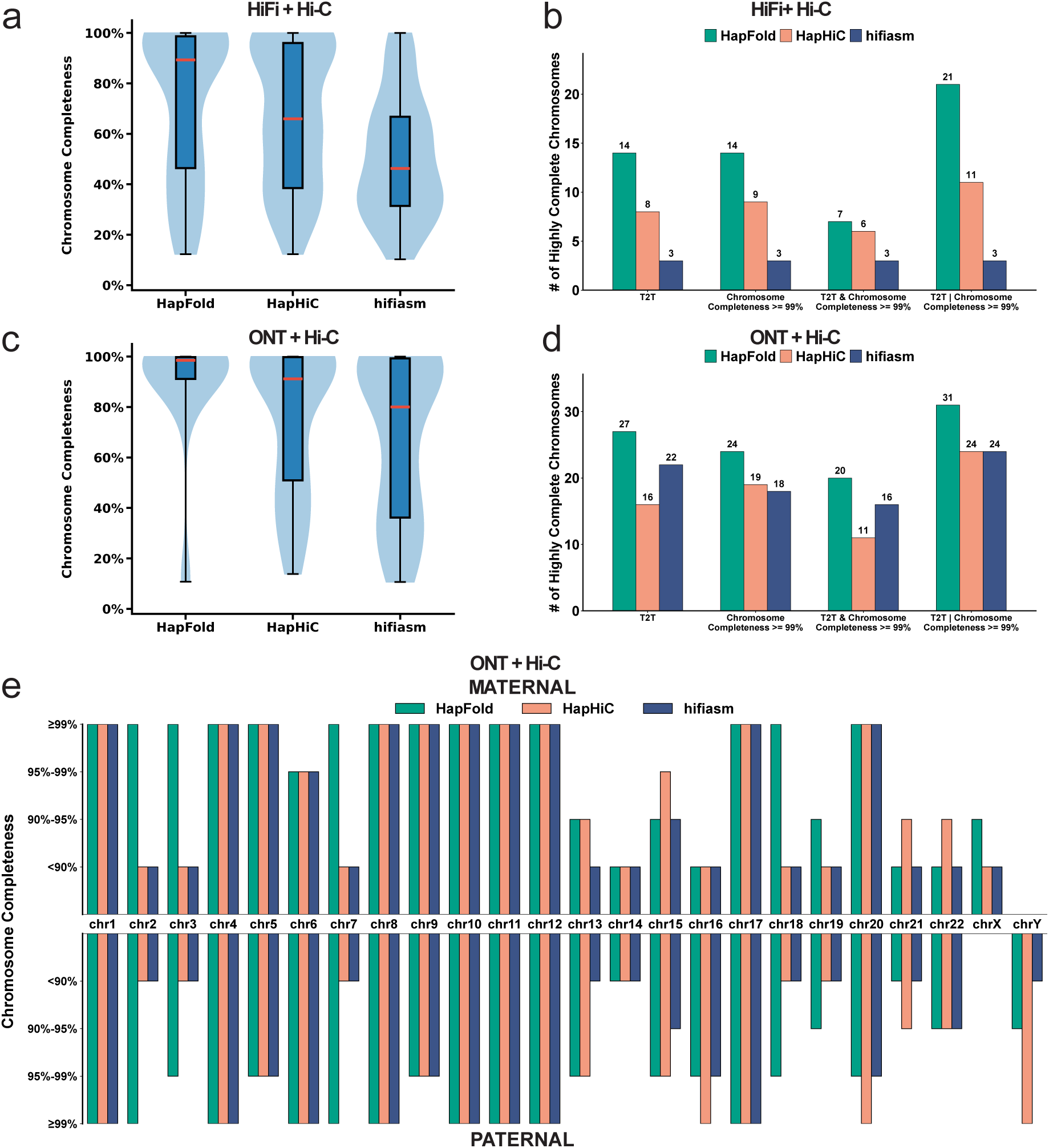
Evaluation of HG002 genome assembly T2T-level and completeness. **a**, Chromosomal completeness distributions for assemblies generated using HiFi reads. The violin plots illustrate the completeness ratio (genome fraction) of individual scaffolds relative to their assigned reference chromosomes for the HapFold, HapHiC, and hifiasm assemblies. **b**, # of highly complete chromosomes within the HiFi dataset. A chromosome is classified as “highly complete” if it achieves either T2T-level or ≥ 99% completeness. T2T denotes the count of scaffolds featuring telomeric sequences at both ends (consistent with HPRC identification criteria, potentially containing internal gaps). Completeness ≥ 99% indicates scaffolds covering more than 99% of the corresponding reference chromosome length. The “T2T & Completeness ≥ 99%” category represents scaffolds simultaneously meeting both criteria, whereas “T2T | Completeness ≥ 99%” reflects the total number of chromosomes achieving at least one of these two standards. **c**, Chromosomal completeness of assemblies generated from ONT simplex reads, with distributions plotted as in **a**. **d**, # of highly complete chromosomes within the ONT simplex dataset. The evaluation criteria for “highly complete” chromosomes are identical to those detailed in **b**. **e**, Detailed chromosome-level completeness assessment for the ONT assemblies. Alignment completeness is displayed via a bidirectional vertical bar chart, symmetrically plotting the maternal (upper) and paternal (lower) haplotypes for each chromosome (chr1–chr22, chrX, and chrY). The vertical bars are partitioned to denote four distinct completeness categories (≥ 99%, 95–99%, 90–95%, and *<* 90%) to illustrate the performance of each scaffolding method across the genome.

Using the HiFi dataset, as shown in Fig. 5a, HapFold exhibited a substantial advantage in chromosomal completeness. The median completeness achieved by HapFold represented a nearly twofold increase over hifiasm and outperformed HapHiC by approximately 40%, indicating that scaffolds assembled by HapFold achieve a significantly higher chromosomal completeness ratio.

Fig. 5b summarizes these results using logical intersection (&) and union (|). Specifically, the intersection denotes scaffolds that simultaneously achieve ≥ 99% reconstruction completeness and satisfy the near-T2T criteria by retaining telomeres at both ends, whereas the union encompasses scaffolds meeting at least one of these criteria, defining the number of highly complete chromosomes. In the HiFi dataset, HapFold generated over five times as many near-T2T scaffolds as hifiasm and exceeded HapHiC’s output by approximately 75%. These striking metrics clearly indicate that HapFold possesses a remarkable capability to achieve T2T-level reconstructions utilizing PacBio HiFi sequencing. It is also worth mentioning that even for HapFold, the number of scaffolds satisfying the intersection of both completeness ≥ 99% and near-T2T status was significantly reduced, highlighting the profound biological and algorithmic difficulty of realizing true T2T assemblies. Nevertheless, HapFold—a framework specialized for T2T-level haplotype reconstruction—successfully assembled 21 high-quality chromosomes (out of 46 in the HG002 genome) that met either the near-T2T or high-completeness criteria.

Using the ONT dataset, as shown in Fig. 5c, HapFold achieved exceptional chromosomal completeness. Specifically, it reached a median completeness of approximately 100%, representing a 20-percentage-point increase over hifiasm(Hi-C) (approximately 80%) and surpassing HapHiC (approximately 90%). These observations indicate that leveraging the inherent advantages of ONT over HiFi assemblies, hybrid scaffolding can further elevate chromosomal completeness to optimal levels.

As illustrated in Fig. 5d, HapFold produced the largest number of scaffolds across all evaluated categories. Specifically, HapFold reconstructed 27 T2T scaffolds, surpassing the 22 and 16 scaffolds identified by hifiasm and HapHiC, respectively. This performance advantage extended to sequence completeness, where HapFold resolved 23 scaffolds with ≥ 99% completeness, compared to 18 for hifiasm(Hi-C) and 21 for HapHiC. Under the most stringent criteria—requiring both T2T-level and ≥ 99% completeness—HapFold resolved 19 chromosomes, outperforming both baseline methods (16 for hifiasm and 11 for HapHiC). Notably, HapFold also achieved high-completeness reconstructions for over two-thirds of the chromosomes (32 of 46). These results suggest that integrating HapFold significantly enhances the continuity and completeness of chromosome-scale assemblies.

We further evaluated the maximum chromosomal completeness achieved for each individual chromosome (Fig. 5e). For this metric, we identified the single assembled scaffold that covered the largest proportion of a given reference chromosome. For instance, a completeness of ≥ 99% indicates that a single continuous scaffold successfully spans over 99% of the target chromosome’s total length. Benefiting from ultra-long read lengths and near-HiFi accuracy, ONT R10.4.1 simplex data inherently yields assemblies with superior overall contiguity and structural accuracy compared to HiFi datasets. As illustrated in the results, HapFold exhibits a distinct advantage in achieving a high degree of chromosomal reconstruction. Notably, for both the maternal and paternal haplotypes of chr2, chr3, and chr7, HapFold achieves a completeness of ≥ 99%. In contrast, the corresponding maximum scaffolds from hifiasm and HapHiC fail to exceed the 90% completeness threshold on these chromosomes. This difference indicates HapFold’s ability to resolve complex or repetitive genomic gaps that often fragment sequence-based scaffolders. Beyond these specific chromosomes, a broader look at the results reveals that both HapFold and HapHiC demonstrate varying degrees of improvement over hifiasm’s contigs across multiple chromosomes. This trend suggests that even with highly contiguous ONT simplex data, dedicated scaffolding tools can still be effectively utilized to further enhance chromosomal completeness. Ultimately, as evidenced by the comprehensive evaluations of T2T counts and chromosomal completeness, HapFold is capable of producing a greater number of T2T scaffolds and consistently achieves a higher degree of chromosome-level reconstruction across the majority of the genome.

Finally, to systematically evaluate the assembly differences between PacBio HiFi and standard ONT simplex reads, we compared the assembly gaps within segmental duplications (SDs) and centromeres across the HapFold and hifiasm results aligned to the HG002 reference genome (Extended Data Fig. 5). Within the HapFold assemblies (Extended Data Fig. 5a), we observed that ONT does not universally outperform HiFi data in resolving complex repetitive and centromeric regions. For instance, across specific loci such as the paternal centromere of chromosome 9, HapFold achieved superior contiguity using HiFi reads compared to ONT. Collectively, these findings indicate that while ONT generally yields a higher overall number of T2T-level chromosomes, HiFi sequencing—when empowered by HapFold—can still deliver superior assembly performance in specific genomic regions that remain recalcitrant to ONT profiling.

As depicted in Fig. 5b, HapFold applied to HiFi data delivers T2T-level assembly performance nearly on par with hifiasm using ONT data (21 and 24 highly complete chromosomes, respectively). This structural advantage is vividly illustrated in the Extended Data Fig. 5b. Specifically, across complex SD and centromeric regions where hifiasm assemblies suffer from contig fragmentation, HapFold successfully bridges these gaps to re-establish scaffold continuity. This observation is highly significant, as it provides compelling evidence that when leveraged by advanced scaffolding algorithms like HapFold, HiFi sequencing can successfully overcome its inherent length limitations to achieve exceptional T2T-level assembly outcomes.

Beyond generating highly contiguous individual chromosomes, an overall comparative analysis highlights HapFold’s superior structural resolution over hifiasm. Crucially, while the exact nucleotide sequences within these gaps remain unfilled, HapFold accurately localizes these gaps and links them to form contiguous scaffolds. This establishment of long-range chromosomal connectivity is paramount, as it preserves macro-genomic architecture and provides an essential structural backbone that precisely guides subsequent T2T gap closure.

### Haplotype-resolved reconstruction of complex genomic regions

A critical objective in modern genomics is the accurate reconstruction of structurally complex regions. These challenging loci, frequently embedded within highly repetitive and duplicated genomic architectures, are notoriously difficult to resolve and highly susceptible to scaffolding errors, thereby creating significant bottlenecks for various downstream genomic applications.

Haplotype-resolved whole-genome assembly provides a powerful approach to decipher these intractable sequences. To mitigate potential evaluation artifacts stemming from reference bias, we aligned the assemblies against the highly contiguous HG002 Q100 reference genome, with gene annotations lifted over from the GRCh38 reference assembly. Within this evaluation framework, we directly compared the reconstructive capabilities of HapFold and HapHiC across representative loci characterized by distinct types of complex genomic architectures.

We evaluated regions dominated by short repeats and highly similar alleles, using the Immunoglobulin Heavy Variable (IGHV) gene family as a representative example. IGHV genes are essential components of the adaptive immune system that drive antibody diversity, and they are densely clustered within genomic segments exhibiting extreme allelic sequence homology. Based on the annotated coordinates of the IGHV gene family, we selected two target loci: the paternal chromosomal region at Chr. 14: 105.50–105.53 Mb and the maternal region at Chr. 14: 108.18–108.43 Mb. As illustrated in Figs. 6a and 6b, HapFold successfully and continuously reconstructed these highly homologous gene clusters on both haplotypes. In contrast, HapHiC failed to accurately resolve these loci, exhibiting severe fragmentation and structural misjoins across both the maternal and paternal assemblies.

**Fig. 6.**
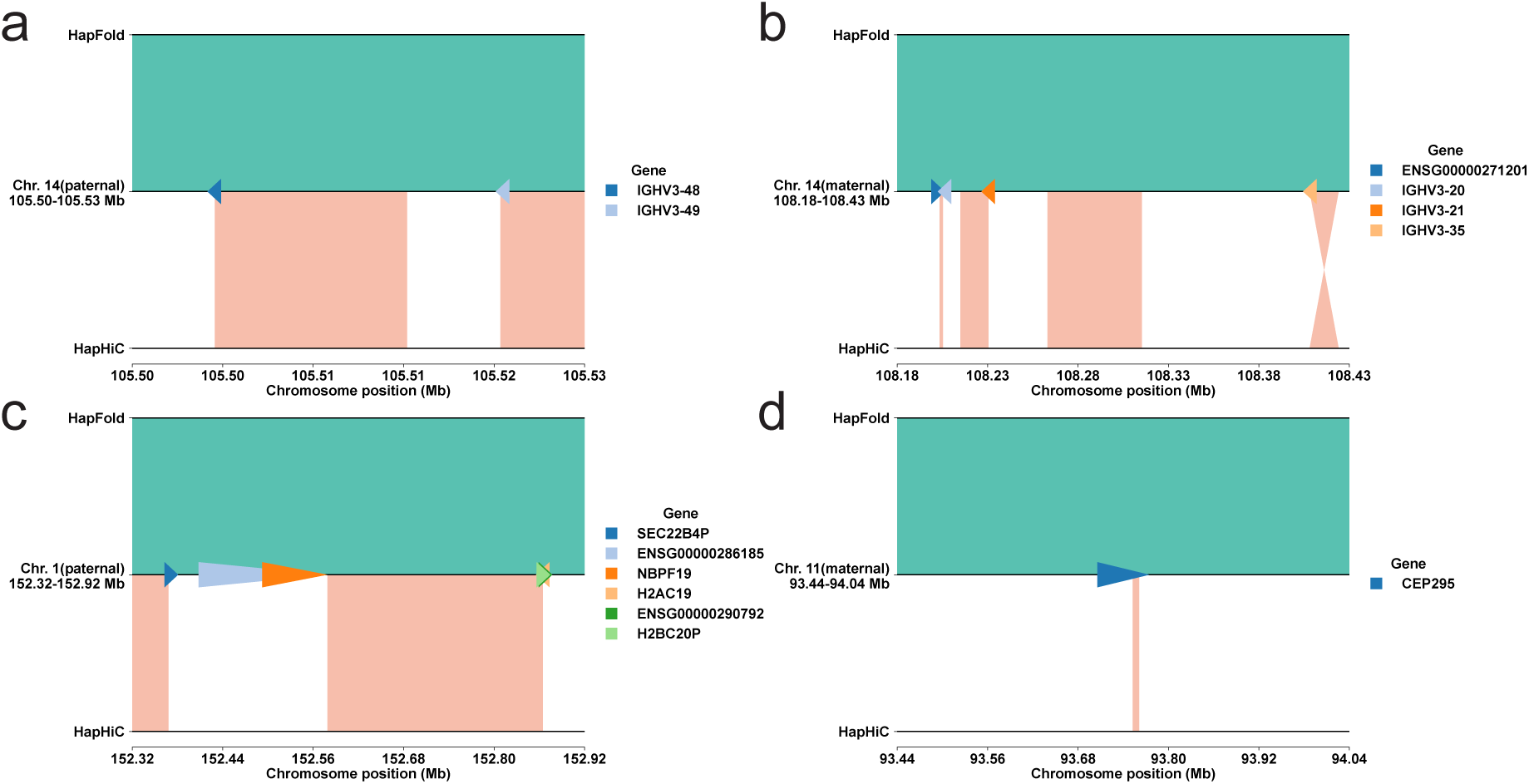
Comparison of HapFold and HapHiC assemblies across complex medically relevant genomic loci in the HG002 genome. Each panel illustrates the alignment of the respective scaffolding assemblies against the HG002 reference genome. The reference axis (central black line) displays annotated gene models as colored arrows, where arrow direction indicates strand orientation and length corresponds to gene size. Alignments generated by HapFold are shown above the reference axis in teal, demonstrating continuous, gap-free sequence reconstruction across these challenging regions. Conversely, alignments generated by HapHiC are shown below the reference axis in salmon, displaying scaffolding fragmentation, structural gaps, and unaligned segments. **a**, Resolution of the highly polymorphic immunoglobulin heavy (IGH) locus on the paternal haplotype of chromosome 14, highlighting the variable region genes *IGHV3-48* and *IGHV3-49*. **b**, Resolution of the corresponding IGH locus on the maternal haplotype of chromosome 14, featuring the variable region genes *IGHV3-20*, *IGHV3-21*, and *IGHV3-35*. **c**, Reconstruction of a highly repetitive segmental duplication hotspot on the paternal haplotype of chromosome 1, encompassing *NBPF19* and adjacent structurally complex genes. **d**, Assembly continuity across the *CEP295* locus on the maternal haplotype of chromosome 11.

Furthermore, we investigated regions characterized by high-homology sequences and extremely repetitive multi-copy domain genes, optimally exemplified by pseudogenes and the Neuroblastoma Breakpoint Family (NBPF) (Fig. 6c). Pseudogenes frequently retain *>* 99% sequence identity with their functional parental counterparts, while NBPF genes feature massive tandem repeat arrays. During scaffolding, these identical segments and extreme copy number variations frequently induce scaffolders to produce structural misassemblies.

As illustrated in Fig. 6c,d, we compared the reconstruction performance of HapFold and HapHiC at two representative complex loci. At the paternal Chr. 1: 152.32–152.92 Mb region, a dense cluster containing the multi-copy gene *NBPF19*, the pseudogene *SEC22B4P*, and other highly similar loci such as *ENSG00000290792* and *ENSG00000286185*, HapFold reconstructed a gapless, contiguous sequence, whereas HapHiC exhibited distinct assembly gaps (Fig. 6c). A consistent pattern was observed at the maternal *CEP295* locus on Chr. 11: 93.44–94.04 Mb, where HapFold preserved local structural continuity, while HapHiC showed localized fragmentation and unspanned gaps (Fig. 6d).

Ultimately, while the initial reconstruction of genomic regions heavily depends on the read-overlap resolution of upstream assemblers, it imposes equally stringent demands on downstream scaffolding frameworks. Guaranteeing high-fidelity, extended scaffold contiguity is an essential prerequisite for the precise identification of complex genes.

### Runtime and memory usage evaluation

To evaluate the computational requirements of HapFold, we benchmarked its runtime and peak memory usage across five datasets of varying genomic complexity (Fig. 7). For the sequence-based scaffolders (YaHS, HapHiC, SALSA2, and 3D-DNA), the input consists of alignment files generated through a standardized BWA-based Hi-C mapping pipeline. In contrast, HapFold and pstools directly utilize their internal graph-aware mapping modules. The total runtime for all scaffolding workflows is decomposed into two stages: Hi-C mapping and the subsequent scaffolding process.

**Fig. 7.**
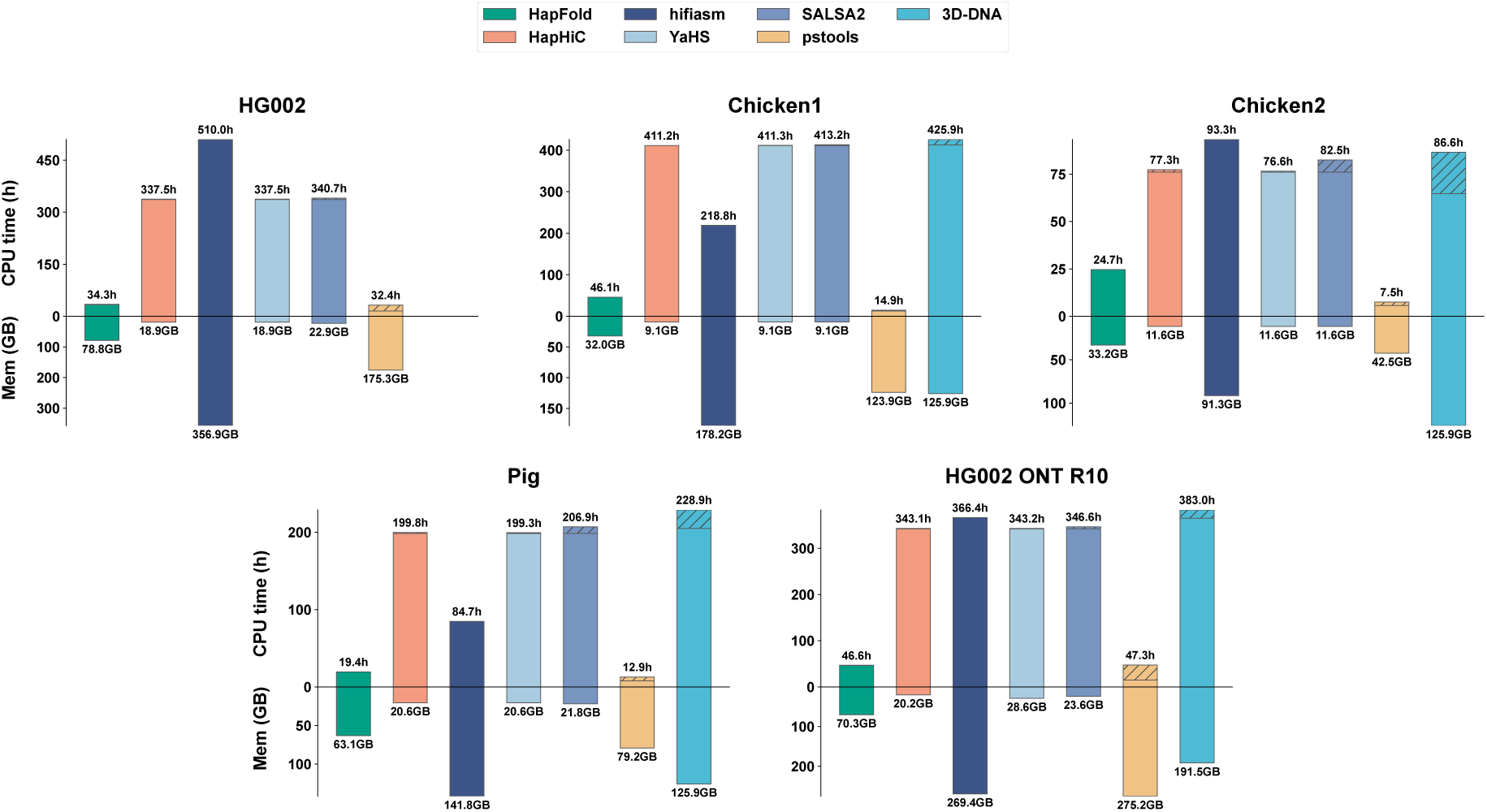
Runtime and peak memory usage of scaffolding tools. The bar charts illustrate total CPU time in hours (positive y-axis) and peak memory usage in GB (negative y-axis). Total CPU time is the sum of the mapping and scaffolding durations, where the hatched portions of the bars specifically denote the time spent on the scaffolding stage. Peak memory represents the maximum memory footprint recorded during the execution of each tool. Note that 3D-DNA failed to complete the assembly on the HG002 dataset due to a bug in its internal script that triggered an infinite loop. Note that computational resource usage for hifiasm is included as a baseline, corresponding to the initial genome assembly step performed before scaffolding.

HapFold demonstrates a substantial runtime advantage, primarily driven by its efficient mapping strategy. Across all benchmarks, HapFold consistently outperforms current sequence-based scaffolders, achieving up to a 10-fold speedup compared to tools such as HapHiC and YaHS. For instance, in the HG002 dataset, HapFold completed the entire workflow in 34.3 hours, whereas the BWA-based pipelines required over 330 hours. Notably, for the HG002 benchmark specifically, 3D-DNA was excluded as its execution proved computationally prohibitive, failing to yield results within a reasonable timeframe. This computational efficiency stems from a *k*-mer based mapping strategy utilized by both HapFold and pstools, which significantly accelerates the alignment process. With the exception of hifiasm, most existing scaffolders (such as HapHiC) rely on external aligners like BWA to perform computationally expensive read-to-contig mapping. By avoiding this bottleneck, HapFold achieves rapid execution times while consistently yielding assemblies of superior structural quality.

Regarding memory consumption, hifiasm serves as a critical reference point, as it is a prerequisite for generating the input unitig graphs. Across all five benchmarks, the peak memory usage of hifiasm is significantly higher than that of any downstream scaffolder, reaching 356.9 GB for the HG002 sample. HapFold maintains a moderate memory footprint, ranging from 32.0 GB to 78.8 GB across the diverse datasets, ensuring its resource requirements are well-supported throughout the entire genome assembly workflow. In comparison, HapHiC, SALSA2, and YaHS exhibit nearly identical memory consumption proportions (e.g., ∼ 18.9 GB for HG002 and ∼ 9.1 GB for Chicken1), which stems from their shared reliance on the same BWA-based alignment pipeline for processing Hi-C data. In contrast, pstools requires substantially higher memory resources, with peak usage reaching 175.3 GB for HG002 and soaring to 275.2 GB for the HG002 ONT R10 dataset. Given that users must already allocate substantial memory for the initial hifiasm assembly, HapFold’s memory usage does not impose an additional practical bottleneck for the overall workflow. In summary, HapFold achieves an order-of-magnitude improvement in execution speed compared to traditional scaffolders while maintaining a reasonable memory footprint. Together, these computational advantages ensure its robust applicability and scalability for large-scale genome assemblies.

## Discussion

The rapid evolution of sequencing technologies and assembly algorithms has the potential to overcome a final hurdle in contemporary genomics. T2T haplotype-resolved assemblies mean the optimum at which one can display the DNA of diploid individuals, both in terms of completeness and evolutionary resolution. Therefore, such assemblies establish a canonical standard in ancestry-aware genomics.

In recognition of its greater meaning, and encouraged by the novel opportunities provided by advances in sequencing technologies, recent assembly approaches have decidedly been pursuing this standard. However, while the new opportunities bring that ultimate goal in sight, most recent approaches have also pointed out the struggle it means to actually reach it: one needs to overcome various seriously challenging technical pitfalls.

Here, we have presented HapFold as an approach that has successfully addressed the remaining concerns. While the remaining issues have been resolved, HapFold has maintained the high standards with respect to earlier, more traditional challenges. Namely, HapFold has delivered a way that eliminates the trade-off between the structural accuracy of the assemblies, which is important to accurately distinguish between the ancestral phases, and their contiguity, which is important to deliver T2T type, full length assemblies. All prior approaches had to trade evolutionary resolution for completeness or vise versa. HapFold is the first approach to computing assemblies that excel in both categories.

HapFold’s key to success has been to synthesize graph-based assembly techniques—which emphasize the structural accuracy, so decisively promote haplotype-level resolution—and sequenced-based techniques, where the latter decisively enhance the contiguity, and in that the completeness of the assemblies. As documented by standard benchmarking experiments, HapFold has proven to be the only approach to deliver T2T haplotype-resolved assemblies that set new standards across the entire range of quality criteria for such assemblies.

In that sense, the wider meaning of HapFold is to successfully unify two paradigms that appear to be contradictory at first glance. While sequence-based protocols stress the sequential order of genomic elements, recently increasingly popular graph-based protocols stress the disentanglement of genetic variation in terms of their assignment to the ancestors of the individual organism under investigation. With both of these computational paradigms having merits in their own right, only very few approaches have attempted to get them under one umbrella. One of the reasons certainly is the fact that the approaches following the more traditional sequence-centric paradigm appeared to have reached their ultimate limits, which made further investments in this paradigm look pointless. In that particular vein, one can interpret HapFold as an approach that revives the older paradigm, by successfully marrying it with the more advanced and currently more popular graph-centric paradigm.

In more detail, HapFold addresses the reference-free, haplotype-aware scaffolding of (already haplotype-aware) contigs as the driving challenge in the context of the usual computational steps when generating chromosome-length, ancestry-aware genome assemblies. The fundamental advantage of HapFold is to integrate the underlying graphs also during scaffolding, and in that to exclusively stitch together contigs from identical ancestors. Since all earlier approaches make use of graphs only when assigning contigs to the two different ancestral phases, this establishes a decisive novelty.

HapFold establishes a critical algorithmic bridge by integrating graph topology during the initial stages in a way that delivers intermediate assemblies that one can connect in an ancestry-aware manner even across the gaps that one remains to scaffold. Guided by these newly equipped, and highly resolved intermediary structures, subsequent application of sequence-based modules allows to pick up the ancestry marks generated in the graph-based assembly stage, which leads to generation of decisively more complete while still ancestry-aware assemblies. In somewhat more technical detail, a dedicated Hidden Markov model (HMM) processes the refined inputs, which enables HapFold to effectively and near-exhaustively resolve the remaining ambiguities. It is important to realize that HapFold is a hybrid approach insofar as it combines most relevant current sequencing technologies, third-generation sequencing (such as ONT and PacBio type reads), and Hi-C reads to successfully implement the ideas just outlined. The final product are precise, full-length chromosome-level, so T2T reconstructed haplotypes.

Our benchmarking experiments demonstrate that HapFold outperforms all existing competitive haplotype-aware T2T scaffolding approaches on diploid genomes across various, diverse species, encompassing human, animal and plant datasets, across the board of approved assembly quality criteria. Most importantly, when processing standard ONT simplex reads coupled with Hi-C data, HapFold successfully resolved a greater number of near-T2T chromosomes and achieved substantially higher per-chromosome completeness than competing methods, which highlights the truly excellent contiguity of its assemblies. Interestingly, HapFold also exclusively leads PacBio HiFi-based assemblies to deliver a number of near-T2T chromosomes that is comparable with that achieved when using ONT simplex data. This is surprising insofar as PacBio HiFi reads are considerably shorter than ONT simplex reads. This particular result points out that HapFold does not depend on the particular length of the input contigs, despite this length being a crucial factor for alternative scaffolding approaches.

Further results demonstrate that the dual-level hybrid architecture of HapFold provides unique capabilities to keep complex gene families intact during reconstruction. This suggests that our hybrid (graph- and sequence-based) scaffolding framework successfully captures biological nuances that are frequently collapsed or lost in traditional proximity-based pipelines. Furthermore, our evaluations using Pore-C data—–which provides higher-order chromatin contact information—–demonstrate that HapFold outperforms existing tools by more effectively capturing complex contact signals.

At the same time, evaluations across different Hi-C sequencing depths highlight HapFold’s remarkable robustness. HapFold consistently achieves optimal performance in comparison to others, regardless of the particular level of depth. Consequently, this indicates that HapFold can also effectively leverage lower-coverage Hi-C data, rendering high-quality diploid assemblies considerably less expensive. In addition to the reductions in sequencing costs, HapFold offers significant practical advantages in terms of computational efficiency. Compared to conventional scaffolders that rely on read-to-contig alignments for Hi-C mapping, HapFold operates approximately an order of magnitude faster. HapFold achieves this acceleration by adopting an efficient *k*-mer-based mapping strategy, particularly designed to process unitig graphs. This effectively eliminates the computational bottlenecks typically associated with complex genome assembly.

Despite the superior performance of HapFold, there are still challenges whose consideration is an obviously worthwhile undertaking. HapFold is currently optimized specifically for diploid genomes and does not yet support the assembly of polyploid species. Extending the hybrid scaffolding framework that is the foundation of HapFold, to also accommodate the higher complexity of polyploid genomes represents a key direction for our future work. Having extended HapFold to also polyploid organisms will mean to be able to compute T2T, so chromosome-level full-length assemblies that resolve the ancestral lineages for every organism for which parent type ancestry matters.

Notably, HapFold consistently achieves higher numbers for near-T2T haplotype-resolved diploid assemblies across multiple sequencing modalities. Using standard ONT simplex reads, it achieves a superior yield of assemblies with ≥ 99% chromosomal completeness. Strikingly, when processing HiFi and Hi-C datasets, HapFold elevates assembly contiguity to yield a comparable number of highly complete and near-T2T chromosomes relative to the latest ONT-based hifiasm results. This dual capability marks a significant step toward low-cost, telomere-to-telomere genomics. By comprehensively resolving allelic diversity and complex structural variations, HapFold is well-positioned to facilitate large-scale pangenome endeavors and advance our understanding of the genetic underpinnings linking complex genomic architecture to phenotypic outcomes and human diseases.

## Methods

### Read mapping

The foundation of the HapFold scaffolding process is the unitig graph, which explicitly represents heterozygous variations as bubble motifs (Fig. 1b). HapFold operates on the unitig as its fundamental computational unit. Structurally, these nodes represent continuous, high-confidence genomic segments formed by perfectly overlapping reads, free from any internal branches or structural ambiguities. The directed edges connecting these nodes encode the sequence overlaps between adjacent unitigs. In these structures, nodes are categorized into homozygous junctions (shared by both haplotypes) and heterozygous unitigs (haplotype-specific paths). Standard scaffolding pipelines typically rely on linear base-level alignment tools, such as BWA [58] or minimap2 [59], to map Hi-C read pairs. However, this conventional approach faces significant challenges when applied to diploid graphs. Specifically, it suffers from severe allelic ambiguity in homozygous or repetitive regions, where reads are frequently misassigned to the incorrect haplotype path due to sequence identity. Furthermore, the computational overhead of base-level alignment becomes prohibitive when processing the deep-coverage proximity data required for chromosome-scale assembly.

To address these limitations, HapFold implements a rapid, graph-aware mapping strategy based on unique *k*-mer indexing. To process the sequence information embedded within the assembly graph, the algorithm first decomposes the constituent unitigs into *k*-mers (default *k* = 31). For each *k*-mer, the tool records its coordinate within the sequence and its corresponding unitig identifier. To optimize efficiency, HapFold stores only the canonical representation of each *k*-mer—defined as the lexicographically smaller string between the forward sequence and its reverse complement—effectively reducing the memory consumption.

These *k*-mers are indexed in a high-performance hash table architecture analogous to those employed in minimap2 and hifiasm [22], ensuring highly efficient storage and retrieval. Given that unitig graphs of diploid genomes contain vast homozygous regions with identical sequences, HapFold applies a rigorous filtering step to ensure mapping specificity. During the indexing stage, the algorithm tracks the occurrence frequency of each *k*-mer across the entire assembly. Any *k*-mer appearing more than once in the graph is flagged as repetitive or homozygous and is subsequently ignored during the mapping process. The resulting index contains only unique *k*-mers, which serve as high-resolution markers for non-homologous or variant-dense regions. This full *k*-mer indexing approach provides superior specificity compared to the minimizer-based methods used by BWA or minimap2, which often sacrifice resolution for smaller index sizes. To further maximize throughput, HapFold employs a pipelined streaming architecture, where sequence retrieval, *k*-mer decomposition, and hash table ingestion are performed concurrently. This design significantly accelerates the overall hash table construction process and ensures that the mapping can scale efficiently with genome size and data depth.

HapFold adopts a streaming architecture to process proximity-ligation data in real-time, where each end of a paired-end read is analyzed independently. Initially, the tool identifies unique *k*-mers within the read and queries the pre-computed hash table to identify potential candidate unitig targets. A consensus orientation is then determined by evaluating the count and polarity (forward vs. reverse-complement) of *k*-mer matches. To ensure high specificity, only *k*-mers consistent with this consensus direction are retained for further validation. Subsequently, the algorithm examines the physical coordinates of these matches to define coherent hit clusters, referred to as anchors. To ensure mapping reliability, HapFold evaluates each candidate based on the total number of anchors and the density of non-overlapping clusters. For the purpose of differentiation, contiguous anchors are treated as a single overlapping anchor; HapFold only considers a unitig as a valid candidate if it contains at least two distinct, non-overlapping anchors. Finally, among all candidate unitigs identified for a Hi-C read, a target is uniquely selected only if it exhibits a significantly superior non-redundant *k*-mer coverage compared to other candidates. In cases of multi-mapping where multiple unitigs present equally plausible hits, the read pair is discarded to maintain the integrity of the proximity signal. To maintain a minimal memory footprint, the algorithm retains only the target unitig identifier and the mapping orientation for each hit; base-level positional information is intentionally discarded at this stage as it is redundant for the subsequent graph-based optimization. After independent mapping, the tool synchronizes the paired-end information. Signal filtering is performed line-by-line: interactions are discarded if either end fails to map or if both ends map to the same unitig. For valid pairs mapping to distinct nodes, the interaction is recorded as a high-confidence link, documenting whether the two unitigs exhibit a forward or reverse-complement orientation relative to one another. The final results are consolidated into a compact four-column format, specifying the two unitig IDs and the aggregate counts of their forward and reverse interaction signals.

Importantly, while existing scaffolding tools necessitate precise base-level coordinates to infer the order and orientation of long contigs, HapFold operates on the unitig as its fundamental computational unit. Because unitigs are formed by unambiguous overlaps of error-corrected reads devoid of any structural branching, they represent exceptionally high-fidelity genomic segments that are inherently much shorter than the final scaffolds. In this framework, unitigs can be integrated as the minimal traversable units, effectively acting as ultra-long synthetic reads. The density and distribution of Hi-C signals anchored to these unitigs provide sufficient resolution for global physical positioning, rendering precise intra-unitig coordinates unnecessary. Furthermore, this strategy inherently bypasses the problem of structural chimeras. Chimeras in conventional assemblies often result from the erroneous concatenation of discordant unitigs into a single linear contig. By scaffolding directly at the unitig level and validating connectivity through both topological overlaps and long-range proximity signals, HapFold ensures that structural continuity is reconstructed from its primary components, eliminating the risk of inheriting or creating chimeric joining errors.

### Unitig-graph resolution

HapFold employs the unphased unitig graph as the primary computational framework for its scaffolding and phasing operations. These graphs are standardly generated by modern assemblers, such as hifiasm, through a stringent string graph construction process. During this initial stage, long high-fidelity sequencing reads (e.g., PacBio HiFi) undergo all-to-all overlap analysis, error correction, and multiple sequence alignment to condense consistent read overlaps into unambiguous segments known as unitigs. Because the unitig graph represents the assembly state directly derived from read alignments, it preserves virtually all original genomic information, capturing intricate biallelic variations and repetitive motifs. Processing this pristine state directly makes it the optimal stage for accurate downstream phasing, avoiding the irreversible information loss frequently seen in aggressively simplified contig graphs.

To prepare the graph for large-scale scaffolding, HapFold deploys a structural curation module dedicated to graph fixing and artifact purging (Fig. 1c). The tool conducts an initial topological scan to assign functional roles to each node based on local connectivity. Nodes are classified into three core types: bubble junctions (the divergence and convergence points of heterozygous alleles), internal bubble nodes (the alternative haplotypic paths), and dangling nodes (nodes with connections in only one direction). HapFold then orchestrates a path-searching and structural expansion algorithm originating from these dangling nodes. Depending on their connectivity profiles and traversal contexts, these nodes are functionally reclassified into endpoints of independent subgraphs, fixable bubble anchors that bridge fragmented heterozygous motifs, or spurious branches marked for immediate pruning.

Following the repair of incomplete bubbles and the excision of erroneous branches, HapFold re-establishes the global connectivity of the graph. The re-established topological connections are strictly categorized into three definitive node types—junctions, internal bubble nodes, and single-path nodes serving as the fundamental operational units for subsequent phasing. Concurrently, HapFold identifies and collapses “fake bubbles”, highlighted in red on the left of (Fig. 1c). These artificial structures are typically induced by localized repetitive elements that forge redundant edge connections between otherwise distinct homologous trajectories. Removing these artifacts is imperative, as their presence would otherwise lead to the erroneous classification of bubble chains and single chains.

With the definitive topology stabilized, HapFold partitions the assembly into distinct genomic chains, categorizing the subgraphs into two primary architectures: bubble chains and single chains. To achieve this structural segregation, the algorithm evaluates two core quantitative metrics: the proportional distribution of classified node types and the read coverage depth of each node, which is extracted directly from the input GFA file. Bubble chains are inherently driven by the biallelic heterozygosity characteristic of diploid genomes. They are definitively identified by a high density of bifurcating bubble structures, coupled with coverage dynamics indicative of homologous loci— specifically, where divergent paths exhibit haploid-level read depth that subsequently re-converges at homozygous anchors displaying diploid-level depth. Conversely, genomic segments characterized by a minimal proportion of internal bubbles and prominent, structurally specific coverage profiles are classified as single chains. Echoing the architectural paradigms established by tools like Verkko [27], these single chains typically correspond to unique biological contexts, including extended homozygous regions, pseudoautosomal regions (PARs), or loci exhibiting extreme heterozygosity. In regions of severe sequence divergence, homologous haplotypes fail to align into a standard diploid locus, traversing the graph as independent, non-pairing paths rather than conventional bubbles. Due to this structural complexity, single chains require an alternative phasing strategy distinct from the approach applied to bubble chains.

As depicted in the “diploid path resolution” module of Fig. 1c, HapFold proceeds to execute structural simplification and pathfinding specifically on the isolated bubble chains. For each established bubble chain, the algorithm computes the absolute in-degree and out-degree of all nodes, deliberately omitting the internal bubble nodes from this calculation. During this abstraction step, the flanking junctions of a bubble treat all internal paths as a singular, consolidated incoming or outgoing edge. This perspective allows HapFold to systematically collapse internal bubble nodes and the boundaries of adjacent minor bubbles. The algorithm subsequently isolates only those nodes where the in-degree and out-degree do not simultaneously equal 1, effectively extracting the critical branch nodes that constitute the graph backbone. This backbone represents a complete, end-to-end path through the graph, explicitly excluding minor tips and localized branches. Meanwhile, the bypassed internal nodes within a complex bubble generate a set of candidate ‘Bubble Paths’ (Fig. 1d). These represent all possible continuous trajectories formed by traversing different combinations of internal nodes from the bubble’s source to its sink. To definitively resolve these hidden structural variants, HapFold defers them to a specialized downstream phasing stage, leveraging spatial Hi-C interaction frequencies to drive a comprehensive two-step resolution: exact intra-bubble path phasing followed by global inter-bubble trajectory matching.

### Haplotype phasing

To accurately resolve the diploid genome, HapFold categorizes the unphased graph structures into two distinct topological classes, applying tailored phasing strategies to each: “single chains” and “bubble chains.” While bubble chains represent canonical paired diploid paths (detailed in Fig. 1d), single chains arise from complex structural divergences and require a dedicated algorithmic approach.

#### Single chains phasing

HapFold implements a specialized graph-based path resolution method for “single chains” (Fig. 1c). This targeted approach is specifically tailored to overcome the significant algorithmic challenges posed by structurally divergent regions, such as pseudoautosomal regions (PARs) and highly heterozygous loci. At these complex genomic coordinates, the unphased unitig graph frequently bifurcates or collapses into shared homozygous anchors, which inherently causes conventional traversal algorithms to terminate prematurely at local sinks.

To resolve these linear trajectories, our algorithm employs a state-aware, length-weighted greedy search algorithm, augmented by a dynamic state relaxation mechanism and spatial Hi-C constraints. For a given single chain subgraph, the path extraction initiates from localized source nodes (e.g., those with an in-degree of zero). At each traversal step, the algorithm prioritizes the extension towards the outgoing edge with the maximum underlying sequence length, ensuring the rapid reconstruction of the most contiguous backbone.

To overcome the limitation where unidirectional constraints of the directed graph force standard path traversals to abruptly terminate at these complex structural nodes, we designed a heuristic path-searching strategy. This strategy accounts for the physical reality that it is precisely this structural anchor and its deeper internal regions that physically bridge the two distinct sequences together. Biologically, these nodes often represent bubble endpoints induced by PARs on sex chromosomes, which mimic the heterozygous bubble structures typically found in autosomes. In standard traversals, visiting such a node strictly locks its reverse complement to prevent cyclic infinite loops, which easily leads to the misassembly of distinct sex chromosomes sharing these highly similar sequences. However, upon encountering these shared structural anchors that retain unvisited predecessor branches, HapFold explicitly overrides the mutual exclusivity constraint. Through this dynamic state relaxation strategy, coupled with localized search depth boundaries to prevent combinatorial explosion, the haplotypic path can successfully thread through the collapsed homozygous region without being prematurely blocked. Although the aforementioned approach can traverse haplotypic paths across collapsed regions, it remains susceptible to erroneous assemblies that mistakenly join distinct chromosomal segments due to these high-similarity regions. Therefore, when encountering such structural ambiguities or when evaluating whether to structurally reverse, connect, or terminate at complex endpoints, the algorithm transitions to a probabilistic evaluation stage bridged by Hi-C signal intensities. To seamlessly stitch discontinuous segments across unresolvable gaps or validate a specific branching decision, we define a spatial linkage score between the current terminating node *u* and a prospective downstream node *v*. The algorithm selects the optimal continuation by determining the candidate node *v* that maximizes

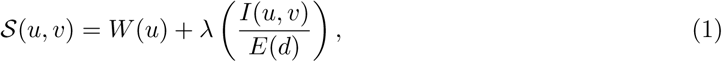

where *W* (*u*) represents the intrinsic Hi-C weight of the current node *u*, establishing a baseline spatial confidence for its extension. *I*(*u, v*) denotes the observed Hi-C interaction frequency between nodes *u* and *v*, *E*(*d*) is the expected background contact frequency at a genomic distance *d* (estimated via shortest topological paths or default gap penalties), and the parameter *λ* serves as a calibrated scaling coefficient to balance the magnitude discrepancy between the nodal weight and the observed-to-expected ratio.

In this formulation, the score S(*u, v*) functions as an absolute topological gateway by evaluating it against a dynamic threshold *τ_u_*, which is derived from the average Hi-C signal of the current node *u*. A resulting score of S(*u, v*) *< τ_u_* implies that the spatial evidence fails to meet the node’s baseline requirement, meaning the Hi-C signal is strictly insufficient to bridge the gap, and the path safely terminates. For candidates yielding a valid passing score (S(*u, v*) ≥ *τ_u_*), HapFold systematically compares them and greedily selects the optimal node *v* that maximizes S(*u, v*). Utilizing this robust, node-specific evaluation threshold, HapFold successfully links genuine haplotypic trajectories while preventing spurious, similarity-driven misassemblies.

#### Bubble chains phasing

Within the graph backbone, the bypassed internal nodes form localized subgraphs comprising multiple alternative contiguous trajectories, termed “bubble paths” (Fig. 1d). To achieve accurate diploid resolution, HapFold executes a spatial-signal-driven phasing strategy. The first critical stage of this process focuses on the intra-chain resolution, specifically the mutual linkage and optimal matching of these internal bubble paths using Hi-C contact frequencies.

For a given genomic interval defined by sequential homozygous anchors, let *B* represent the set of internal bubbles and *P_i_* denote the set of candidate paths within bubble *b_i_* ∈ *B*. To determine the correct haplotypic configuration, the algorithm establishes a multidimensional linkage matrix to aggregate the raw Hi-C spatial signals. For any path *p* ∈ *P_i_*, let *V* (*p*) denotes the set of constituent nodes comprising the path. For any two paths *p* ∈ *P_i_* and *q* ∈ *P_j_* from distinct bubbles (*i* = *j*), HapFold computes the cumulative Hi-C interaction count *C*(*p, q*) by systematically summing the observed interactions between their constituent nodes, given by

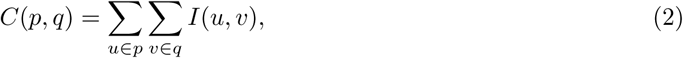

where *I*(*u, v*) denotes the observed Hi-C interaction frequency between nodes *u* and *v*. This quantitative aggregation captures the spatial co-occurrence probability of all possible path combinations.

To resolve the phase from this complex linkage matrix, we implement a greedy, maximum-linkage (Max-link) seed-and-extend algorithm, as conceptually depicted in the central panel of Fig. 1d. The resolution initiates with a seeding stage, where the algorithm scans the entire linkage matrix to identify the global maximum interaction count between any two paths from distinct bubbles. The initial structural seed for the first haplotype trajectory, denoted as the path pair (*p*^∗^*, q*^∗^), is identified by

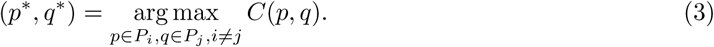

The two most strongly linked paths are therefore confidently assigned to the same haplotype thread. For typical biallelic bubbles, the reciprocal allele paths not selected for this thread are simultaneously assigned to the counterpart haplotype thread, thereby enforcing mutual exclusivity between the two haplotypes.

Following the seeding step, the algorithm enters an iterative topological extension stage. Let *H*_1_ represent the actively growing set of paths assigned to the primary haplotype. In each iteration, HapFold evaluates all currently unassigned bubbles against *H*_1_. For each candidate path *p_k_* ∈ *P_k_* of an unassigned bubble *b_k_*, it dynamically calculates a cumulative extension score *G*(*p_k_, H*_1_), defined as

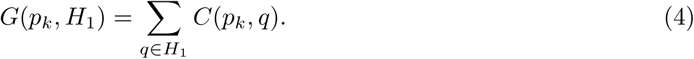

The specific bubble path that exhibits the highest cumulative extension score is iteratively recruited into the primary haplotype set *H*_1_. For typical biallelic bubbles, once an allele path is assigned to *H*_1_, its reciprocal allele path is simultaneously assigned to the counterpart haplotype set *H*_2_, ensuring mutual exclusivity between homologous haplotypes. This greedy extension process continues until all bubbles within the local subgraph are confidently assigned. A complementary extension process is then performed to reconstruct *H*_2_, using only the remaining unassigned paths that are mutually consistent with the established haplotype partition.

This bubble-chain phasing procedure is illustrated in Extended Data Fig. 6. Briefly, a local unitig graph containing multiple bubbles is first refined into a bubble chain, in which the begin and end nodes define the chain boundary and the internal bubble paths retain the alternative haplotype-specific trajectories. Hi-C or Pore-C chromatin contact signals are then used to phase bubble paths both within individual bubbles and across adjacent bubbles. Upon complete partitioning of the internal bubble paths into two mutually consistent haplotype sets, HapFold traverses the bubble-chain backbone and threads through the assigned paths to extract contiguous, haplotype-resolved sequences. Consequently, the entire bubble chain is resolved into one or more pairs of phased contig sequences, as depicted in Fig. 1e. These elongated sequences subsequently trigger the hierarchical scaffolding module, initiating with the intra-chain scaffolding between adjacent subgraphs.

### Scaffolding

Upon completing the localized phasing of all bubble paths, the pipeline transitions to the spatial assembly of these locally phased contigs. To achieve chromosome-scale continuity while maintaining exceptional structural fidelity, HapFold employs a hybrid hierarchical scaffold architecture (Fig. 1e). This framework is structurally divided into two synergistic stages: Local Scaffolding, which relies heavily on graph-based topological traversal, and Global Scaffolding, which transitions to a direct sequence representation and utilizes a dedicated sequence model to finalize the chromosomal architecture.

#### Local Scaffolding

Within each specific subgraph, HapFold performs graph-based scaffolding to stitch the contig pairs into contiguous chromosomal backbones. This local scaffolding process is summarized in Fig. 1e, in which the left panels show the phased subgraphs obtained after bubble-chain phasing, the middle panels show the construction of the primary haplotype-resolved backbone, and the right panels show the reintegration of additional contig fragments guided by chromatin-contact signals. The scaffolding algorithm designates the longest phased contig pair as the initial structural seed. From these anchoring sequences, the algorithm executes a bidirectional topological search, both forward and backward, targeting the adjacent endpoints of neighboring phased contigs. This step corresponds to the transition from the phased local subgraphs to the primary contiguous backbone in Fig. 1e. During this traversal, HapFold encounters two primary connection scenarios. Crucially, any attempt to connect adjacent contigs is strictly predicated on the supporting strength of chromatin-contact signals. If the identified adjacent contigs share a valid, traversable path within the unitig graph, they are seamlessly merged based on their exact sequence overlaps. Conversely, if the adjacent contigs are topologically unreachable in the DAG due to complex local tangles or missing edges, they are linearly concatenated and bridged by inferred gaps. The length of these gaps is inferred from alternative sequences spanning the same region. When this is not possible, a default fixed length is applied. Representative local-scaffolding configurations are further illustrated in Extended Data Fig. 7, which shows how phased contigs can be joined through graph-supported overlaps, inferred gaps and chromatin-contact-guided links while maintaining haplotype consistency.

A representative genomic example is provided by a complex tangled region on chromosome 9, in which the two haplotypes exhibit distinct local graph structures (Extended Data Fig. 8a). Within this locus, the paternal haplotype successfully traverses the region as a continuous path, whereas the maternal trajectory forms a bifurcated structure. While the initial phasing stage naturally breaks the assembly at this ambiguous junction, HapFold leverages the global path topology during its scaffolding stage to bridge the fragmented region by introducing an inferred gap. As evidenced by the alignment landscape in Extended Data Fig. 8b, this graph-based scaffolding strategy enables HapFold to uniquely and successfully link the two flanking contigs. In contrast, both the graph-based assembler (hifiasm) and the sequence-based scaffolder (HapHiC) fail to connect this region. Such structural fragmentation in competing tools significantly exacerbates the difficulty of achieving complete downstream T2T genome assemblies.

To ensure the structural fidelity of these contiguous assemblies, HapFold implements a dual-validation mechanism. Primarily, the exact linkage and relative ordering between any two contig end-points are quantitatively determined by the strength of their spatial Hi-C interaction signals. A strong Hi-C contact frequency dictates the scaffolding confidence and guides the connections across the gaps. Furthermore, HapFold utilizes the phased hap contigs generated by hifiasm as an auxiliary structural reference. The ordering information of the constituent unitig sequences within these contigs assists HapFold in rectifying erroneous orientations and validating the structural continuity of the scaffold.

Once these primary contiguous backbones are established, HapFold initiates a targeted reintegration step to rescue small orphaned contig fragments that were disconnected during the initial path search. Because the constructed scaffold strictly retains the positional coordinates of its constituent contigs, the algorithm can dynamically ‘open’ the sequence at specific loci to accommodate these structural insertions. The viability of anchoring a small contig at a candidate position is rigorously verified using remote connection signals. Furthermore, strand-specific contact frequencies strictly dictate the precise relative ordering and forward/reverse orientation of these fragments during integration. Ultimately, this local scaffolding stage successfully stitches the fragmented sequences, yielding at least one pair of maximally extended phased sequence backbones for each subgraph.

#### Global Scaffolding

Following the construction of the primary phased backbones, HapFold shifts its algorithmic paradigm from abstract graph topology to a direct sequence representation. To maximize assembly completeness and orchestrate macro-level connections, this stage employs a dedicated sequence model, specifically a Hidden Markov Model (HMM) framework, alongside global spatial agglomeration.

Initially, subgraphs often retain unassigned small contigs that were bypassed during the local graph traversal due to topological discontinuities. To anchor these orphaned sequences, HapFold models the placement process analogously to decoding in an HMM. The established sequence backbone serves as an ordered sequence of spatial hidden states (docking loci), while the unassigned small contigs act as independent observations. Every constituent node *w* on the backbone strictly retains its exact genomic coordinate *x*(*w*) established during initial path generation, functioning as a quantifiable docking port.

Let *S* denote the set of unassigned small contigs partitioned by haplotype identifiers, *V* be the set of phased contigs comprising the corresponding backbone, and N (*c*) denote the set of constituent unitigs (internal nodes) within any given contig *c*. To determine the optimal insertion sequence, the algorithm evaluates the spatial “emission” probability for each pair (*s, v*), where *s* ∈ *S* and *v* ∈ *V*, by constructing a multidimensional orientation-aware connectivity matrix. For any small contig *s* and backbone contig *v*, HapFold computes a distance-constrained interaction score *F* (*s, v, o_s_, o_v_*) across all four combinations of relative orientations *o_s_, o_v_* ∈ {0, 1}, representing forward and reverse strands. The raw interaction is aggregated by

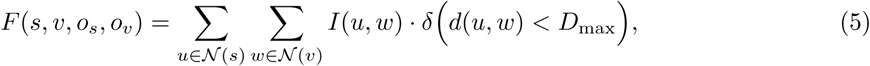

where *I*(*u, w*), as defined above, denotes the observed Hi-C contact frequency between nodes *u* and *w*, and *d*(*u, w*) represents the estimated relative linear distance between these internal nodes based on their strand-specific terminal coordinates. The indicator function *δ* filters out topological noise by rejecting interactions that exceed a maximal spatial threshold *D*_max_, which is dynamically calibrated based on the median length of the assembled contigs within the dataset.

To mitigate length bias, HapFold applies a normalization step. The final objective linkage score L is defined as

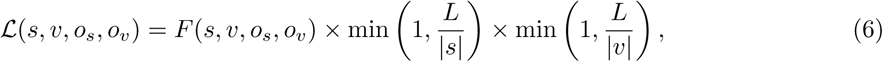

where *L* represents an empirically derived scaling threshold, and |*s*| and |*v*| denote the respective sequence lengths. All valid candidate insertions with L *>* 0 are ranked in descending order. HapFold employs a global greedy optimization strategy, iteratively selecting the highest-scoring candidate and determining the precise insertion locus based on optimal orientation states. To prevent coordinate collisions, multiple insertions into the same backbone are processed in a spatially descending order (from the 3’ end to the 5’ end), dynamically recalculating the global coordinates *x*(*u*) upon every successful integration.

Finally, to bridge the remaining gaps and reach the target chromosomal architecture, the algorithm utilizes the expected haploid chromosome count as a target parameter. If the current number of scaffolds exceeds this biological threshold, HapFold initiates a global sequence-based agglomeration. The algorithm computes a global Hi-C spatial interaction matrix between all available scaffolds. Employing a length-ascending greedy strategy, it systematically prioritizes the shortest scaffolds for integration. For each iteration, HapFold identifies the optimal pairing candidate exhibiting the maximum spatial interaction score. This score is rigorously evaluated against a dynamically calibrated threshold. If the interaction strength securely surpasses this minimum signal requirement, the sequences are concatenated; otherwise, the hypothetical join is aborted. This iterative global merging continues until the total scaffold count converges to the target chromosome number, ultimately yielding a near chromosome-level, haplotype-resolved genome assembly.

#### Handling of ribosomal DNA (rDNA) arrays

A significant challenge in T2T assembly is the presence of high-copy-number ribosomal DNA (rDNA) arrays. As documented in the development of Verkko2 [60], these regions frequently manifest as complex, multi-chromosomal tangles within the assembly graph due to near-perfect sequence identity across distinct genomic loci (Fig. 1). These tangles not only obscure the underlying graph topology but also severely compromise the reliability of Hi-C data; the extreme homology facilitates massive cross-mapping of Hi-C read pairs, creating spurious proximity signals that conventional scaffolding tools—which often ignore the graph context— mistakenly interpret as evidence for inter-chromosomal linkage.

To address this, HapFold implements specialized safeguards during both the phasing and scaffolding stages. First, during the graph-refining process (Fig. 1c), HapFold identifies signature tangle patterns to mark affected genomic chains as “rDNA-associated.” During the subsequent phasing stages, the algorithm applies increased stringency when evaluating path connectivity within these marked regions. When encountering high topological complexity or fragmented structures, the phaser prioritizes path truncation (breaking the sequence) over risky extensions, thereby preempting the formation of multichromosomal chimeras.

In the scaffolding stages, backbone contigs originating from these marked rDNA chains undergo a rigorous “edge-gradient validation.” Instead of relying on aggregate contact counts, HapFold analyzes the decay of Hi-C signals extending from the contig boundaries. A join is finalized only if a robust interaction gradient is detected, ensuring that the signal significantly exceeds the background noise induced by rDNA-related cross-mapping. By leveraging graph-level intelligence to intercept entanglement signals early, HapFold avoids the catastrophic misassemblies common in existing tools that rely solely on linear sequence alignments in repeat-rich regions.

### Quantitative assessment of scaffolding structural integrity via Gini impurity

To quantify the performance of chromosome-scale scaffolding in terms of topological consistency, haplotype purity, and intra-chromosomal collinearity, we developed a length-weighted evaluation metric based on the Gini impurity framework. This metric utilizes the alignment of assembly scaffolds against a high-quality diploid reference to characterize the degree of structural fragmentation and chimeric errors across the physical topological paths of each chromosome. The fundamental purity metric is derived from the Gini impurity formula, which measures the degree of composition diversity within a single scaffold:

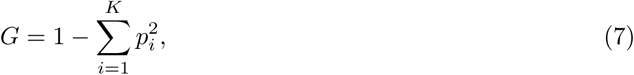

where *K* represents the total number of distinct categorical components (e.g., specific haplotypes or chromosomal origins) identified within the scaffold, and *p_i_* denotes the proportion of the *i*-th component’s aligned length relative to the total aligned length of the scaffold. The Gini impurity was selected for this assessment because it provides a sensitive and bounded measure of assembly fidelity. In an ideal T2T diploid assembly, a scaffold should consist entirely of a single haplotype from a single chromosome, resulting in a Gini impurity of 0. Conversely, structural misassemblies—such as cross-chromosome translocations or inter-homolog phase switches—introduce heterogeneous components that significantly increase the *G* value. By weighting these components by their alignment length, the Gini impurity characterizes structural integrity based on the physical scale of misassemblies. This approach differs from metrics that solely count the frequency of individual misassembly events, thereby ensuring that large-scale structural errors introduced during the scaffolding process are not overlooked due to their potentially low occurrence counts.

To comprehensively evaluate diverse aspects of assembly quality, we implemented this framework through three specific metrics: the Haplotype Gini Impurity (HGI), the Chromosome Gini Impurity (CGI), and the Intra-chromosomal Gini Impurity (IGI). For the calculation of HGI, *K* denotes the parental haplotype identities (i.e., maternal or paternal), where an elevated value reflects the degree of phasing errors within a scaffold. In the case of CGI, *K* represents the distinct primary chromosome identities assigned to the scaffold segments; thus, a higher CGI signifies a greater degree of inter-chromosomal mixing and translocation errors. Regarding the IGI, *K* characterizes the categorical distribution of structural error types relative to the primary chromosome. By assessing the length-weighted proportions of these internal segments—specifically accounting for three distinct types of structural errors—the IGI quantifies the structural integrity within a single chromosome, where an elevated value represents a higher degree of intra-chromosomal structural errors.

## Acknowledgements

This study was supported by Yuelushan Laboratory Breeding Program (No. YLS-2025-ZY03024); Hunan Science and Technology Innovation Plan (No. 2025ZYJ003); the National Natural Science Foundation of China (Grant No. 32400506, 62372159, 62425204, U22A2037); the Natural Science Foundation of Hunan Province (Grant No. 2024JJ4008); Fundamental Research Funds for the Central Universities (Grant No. 541109030062); Fundamental and Interdisciplinary Disciplines Breakthrough Plan of the Ministry of Education of China (JYB2025XDXM602).

## Author contributions

Xiao Luo, Yuansheng Liu and Alexander Schönhuth conceived this study. Yuansheng Liu, Yichen Li and Xiao Luo designed the method. Yichen Li implemented the software. Yichen Li, Yuansheng Liu, Jialu Xu, Zhongzheng Tan and Wenhai Zhang conducted the data analysis. Long Wang, Luohao Xu, Jiawei Luo and Xiangxiang Zeng provided critical feedback and assisted with data interpretation. All authors contributed to writing and revising the manuscript. All authors read and approved the final version of the manuscript.

## Data availability

The datasets supporting the findings of this study are available in public repositories as follows: for the human HG002 genome, the T2T-HG002 v1.1 reference genome was obtained from https://github.com/marbl/HG002 or via NCBI Assembly Accession GCA 018852605.3 (Paternal) and GCA 021951015.3 (Maternal); Hi-C reads at https://s3-us-west-2.amazonaws.com/human-pangenomics/working/HPRC_PLUS/HG002/raw_data/Hi-C/HG002.HiC_1_NovaSeq_1_S1_L002_R1_001.fastq.gz,https://s3-us-west-2.amazonaws.com/human-pangenomics/working/HPRC_PLUS/HG002/raw_data/Hi-C/HG002.HiC_1_NovaSeq_1_S1_L002_R2_001.fastq.gz,https://s3-us-west-2.amazonaws.com/human-pangenomics/working/HPRC_PLUS/HG002/raw_data/Hi-C/HG002.HiC_1_NovaSeq_2_S2_L001_R1_001.fastq.gz,https://s3-us-west-2.amazonaws.com/human-angenomics/working/HPRC_PLUS/HG002/raw_data/Hi-C/HG002.HiC_1_NovaSeq_2_S2_L001_R2_001.fastq.gz, https://s3-us-west-2.amazonaws.com/working/HPRC_PLUS/HG002/raw_data/Hi-C/HG002.HiC_2_NovaSeq_rep1_run2_S1_L001_R1_001.fastq.gz, and https://s3-us-west-2.amazonaws.com/working/HPRC_PLUS/HG002/raw_data/Hi-C/HG002.HiC_2_NovaSeq_rep1_run2_S1_L001_R2_001.fastq.gz; PacBio HiFi reads at https://s3-us-west-2.amazonaws.com/human-pangenomics/working/HPRC_PLUS/HG002/raw_data/PacBio_HiFi/20kb/m64011_190830_220126.Q20.fastq, https://s3-us-west-2.amazonaws.com/human-pangenomics/working/HPRC_PLUS/HG002/raw_data/PacBio_HiFi/20kb/m64011_190901_095311.Q20.fastq, https://s3-us-west-2.amazonaws.com/human-pangenomics/working/HPRC_PLUS/HG002/raw_data/PacBio_HiFi/15kb/m64012_190920_173625.Q20.fastq, https://s3-us-west-2.amazonaws.com/human-pangenomics/working/HPRC_PLUS/HG002/raw_data/PacBio_HiFi/15kb/m64012_190921_234837.Q20.fastq, https://s3-us-west-2.amazonaws.com/human-pangenomics/T2T/scratch/HG002/sequencing/hifi/m64012_201024_012808_Q99-100.reads.fastq.gz, and https://s3-us-west-2.amazonaws.com/human-pangenomics/T2T/scratch/HG002/sequencing/hifi/m64012_201025_074858_Q99-100.reads.fastq.gz; Oxford Nanopore ultra-long (SUP) reads at s3://ont-open-data/giab2025.01/basecalling/sup/HG002/PAW70337; the chicken1 dataset consists of HiFi and Pore-C reads derived from a diploid chicken sample, with its associated sequencing data archived under NCBI BioProject PRJNA1114433; for the chicken2 (*Gallus gallus*, bGalGal1) assembly, Hi-C reads at https://genomeark.s3.amazonaws.com/species/Gallus_gallus/bGalGal1/genomic_data/arima/bGalGal1_S1_L006_R1_001.fastq.gz and https://genomeark.s3.amazonaws.com/species/Gallus_gallus/bGalGal1/genomic_data/arima/bGalGal1_S1_L006_R2_001.fastq.gz; chicken2 HiFi reads at https://genomeark.s3.amazonaws.com/species/Gallus_gallus/bGalGal1/genomic_data/pacbio_hifi/m54306Ue_220613_162445.hifi_reads.fastq.gz, https://genomeark.s3.amazonaws.com/species/Gallus_gallus/bGalGal1/genomic_data/pacbio_hifi/m64055e_220721_053659.hifi_reads.fastq.gz, https://genomeark.s3.amazonaws.com/species/Gallus_gallus/bGalGal1/genomic_data/pacbio_hifi/m64055e_220722_163330.hifi_reads.fastq.gz, and https://genomeark.s3.amazonaws.com/species/Gallus_gallus/bGalGal1/genomic_data/pacbio_hifi/m64330e_220920_184402.hifi_reads.fastq.gz; chicken2 ONT ultra-long reads at https://genomeark.s3.amazonaws.com/species/Gallus_gallus/bGalGal1/genomic_data/ont/ultralong/raw/uncorrected/bGalGal1_PAK10959_20220928.raw.fastq.gz and https://genomeark.s3.amazonaws.com/species/Gallus_gallus/bGalGal1/genomic_data/ont/ultralong/raw/uncorrected/bGalGal1_PAK14466_20220726.raw.fastq.gz; and the pig (*Sus scrofa*) reference genome is hosted on Figshare at https://doi.org/10.6084/m9.figshare.28246343.v1, with associated sequencing data archived under NCBI BioProject PRJNA1139427 (https://www.ncbi.nlm.nih.gov/bioproject/PRJNA1139427); reference genomes for various *Oryza sativa* cultivars were retrieved from the Riceome database at https://riceome.hzau.edu.cn/download/128077.fasta, https://riceome.hzau.edu.cn/download/132278.fasta, https://riceome.hzau.edu.cn/downlo ad/Azucena.fasta, https://riceome.hzau.edu.cn/download/MH63RS3.fasta, https://riceome.hzau.edu.cn/download/127742.fasta, https://riceome.hzau.edu.cn/download/IR64.fasta, https://riceome.hzau.edu.cn/download/127518.fasta, https://riceome.hzau.edu.cn/download/117534.fasta, and https://riceome.hzau.edu.cn/download/117425.fasta;

## Code availability

The source code of HapFold is GPL-3.0 licensed, and publicly available at https://github.com/Luo Group2023/HapFold.

## Declarations

### Ethics approval and consent to participate

Not applicable.

### Consent for publication

Not applicable.

---

**Extended Data Fig. 1.**
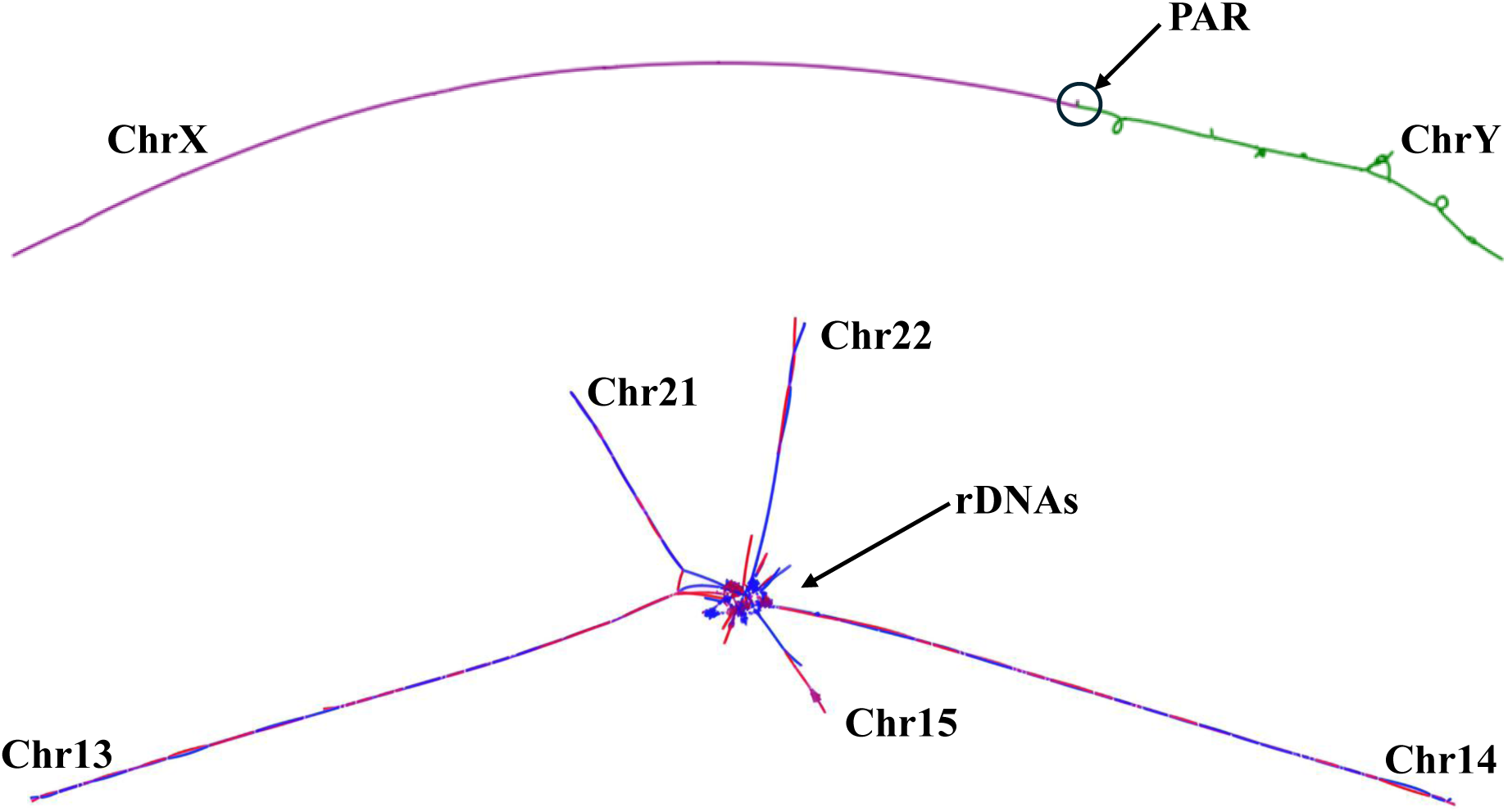
Visualization of complex repetitive regions in the HG002 assembly graph. The top portion of the graph illustrates the linkage between sex chromosomes X and Y mediated by pseudo-autosomal regions (PARs). These regions share high sequence homology, which prevents the sex chromosomes from being resolved as independent components and results in a shared subgraph structure. Below, complex inter-chromosomal connections are observed involving the acrocentric chromosomes 13, 14, 15, 21, and 22. These tangles are driven by the presence of large clusters of ribosomal DNA (rDNA), which possess high sequence identity and a repetitive nature across multiple chromosomes, manifesting as a central tangled hub in the Bandage visualization of the unitig.gfa.

**Extended Data Fig. 2.**
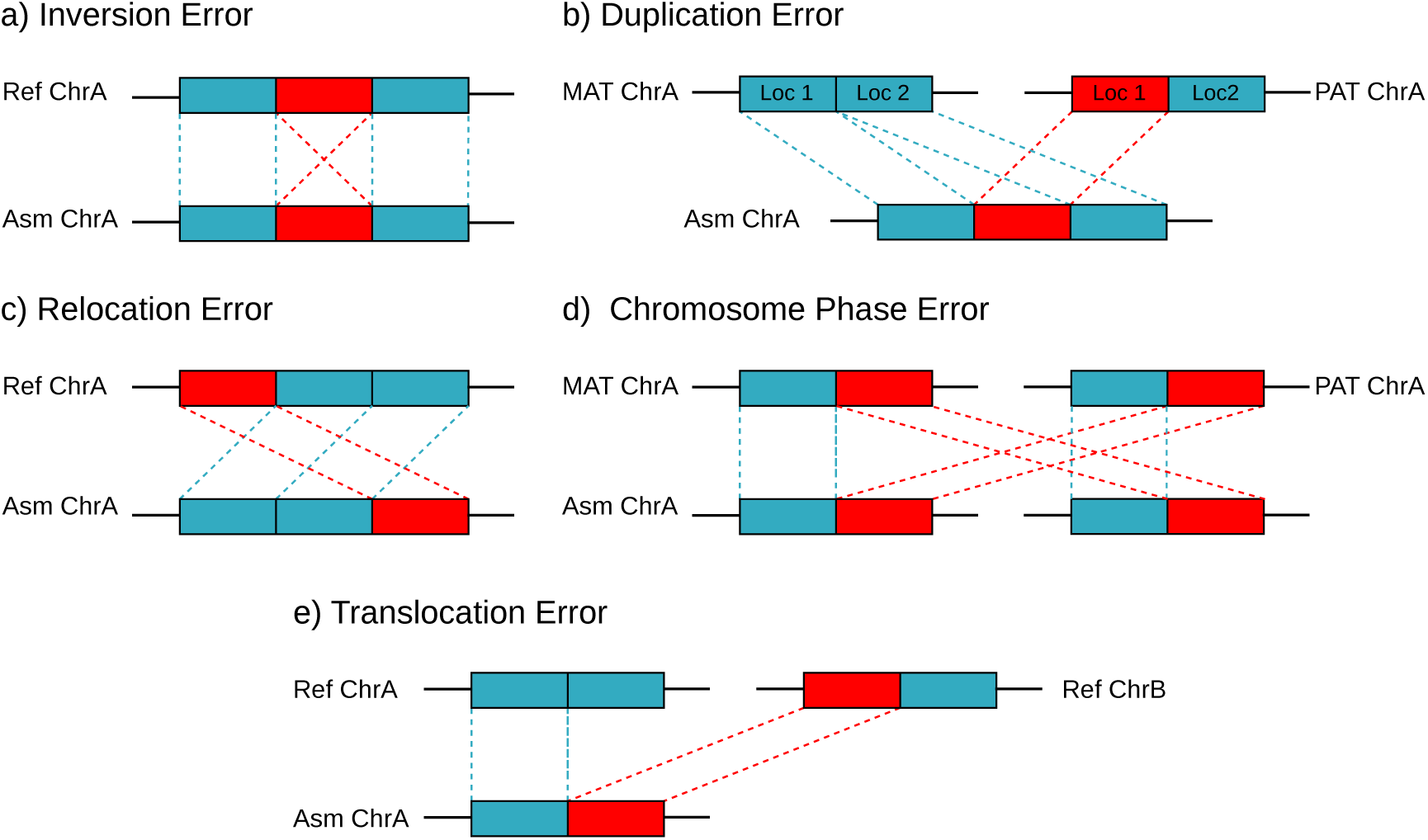
Classification of structural errors in scaffolding assembly. Aligning the assembled sequences (Asm) against the reference (Ref) or the maternal (MAT) and paternal (PAT) haplotypes reveals five primary categories of structural misassemblies. In all panels, red blocks specifically highlight the erroneous segments within the assembly, while dashed lines trace their corresponding sequence alignments. a, Inversion error: a genomic segment is assembled in the reverse physical orientation relative to the reference. b, Duplication error: a genomic region is redundantly assembled, which typically occurs when identical segments from homologous alleles are erroneously concatenated together within the same scaffold. This type of misassembly frequently co-occurs with Chromosome phase errors. c, Relocation error: a contiguous sequence is correctly oriented but misplaced to an incorrect genomic coordinate within the same chromosome. d, Chromosome phase error: an erroneous crossover between maternal and paternal haplotypes occurs within a single scaffold. e, Translocation error: a sequence segment originating from a distinct, non-homologous chromosome (Ref ChrB) is aberrantly fused to the current chromosome (Ref ChrA).

**Extended Data Fig. 3.**
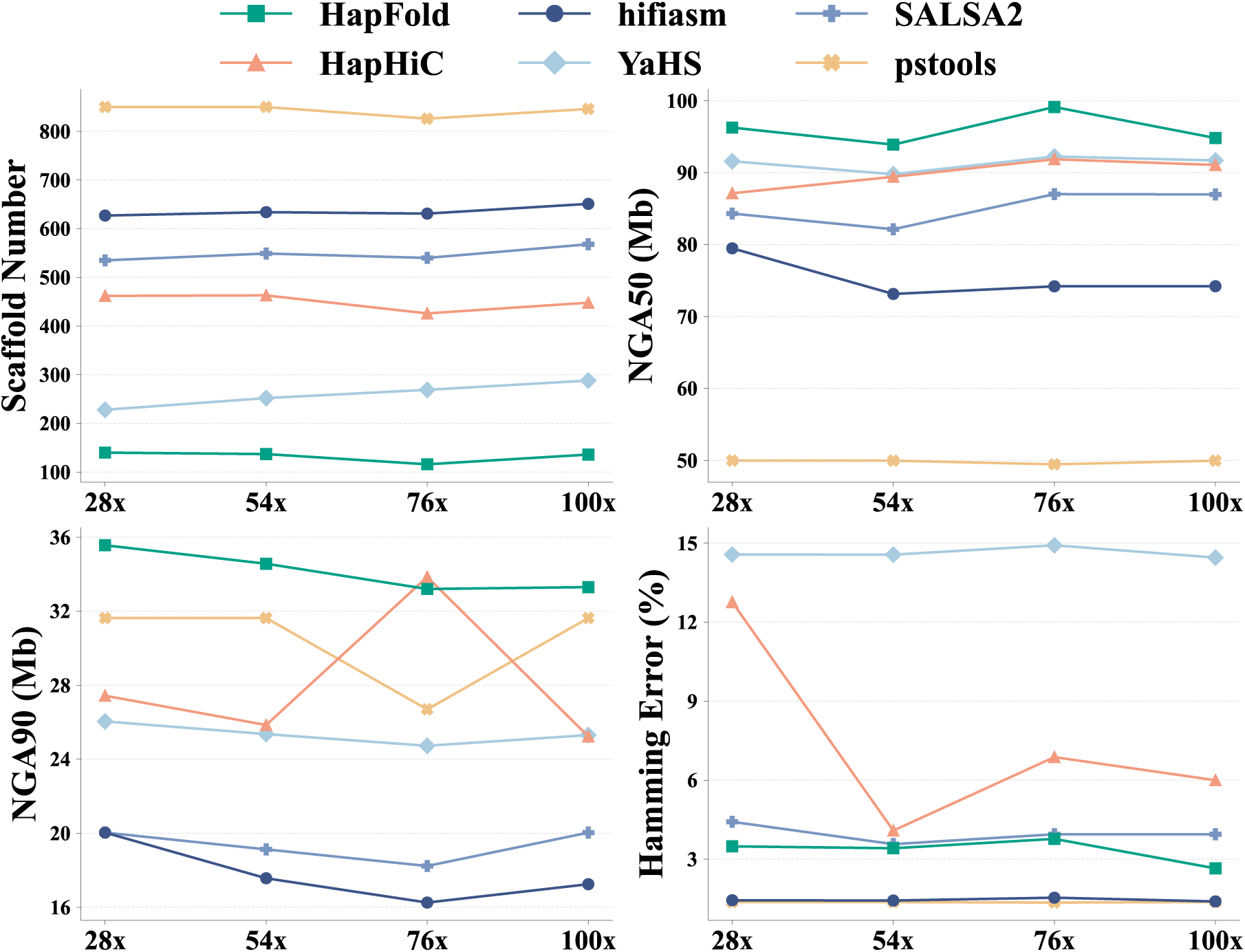
Performance evaluation of scaffolding tools across varying Hi-C coverages on the HG002 dataset. The x-axis denotes the assemblies generated using four different Hi-C sequencing depths (28×, 54×, 76×, and 100×). Scaffold Number indicates the total number of sequences output by each assembly pipeline. NGA50 represents the NG50 length calculated after breaking the scaffolds at structural misassembly breakpoints relative to the reference, reflecting the contiguity of structurally correct blocks. NGA90 follows the same alignment-based penalty but at the 90th percentile, serving as a stringent metric for assessing the structural continuity across the more fragmented and difficult-to-assemble tail ends of the genome. Hamming Error quantifies the phasing accuracy by measuring the proportion of phase switches within the scaffolds, where a lower percentage indicates superior haplotype consistency.

**Extended Data Fig. 4.**
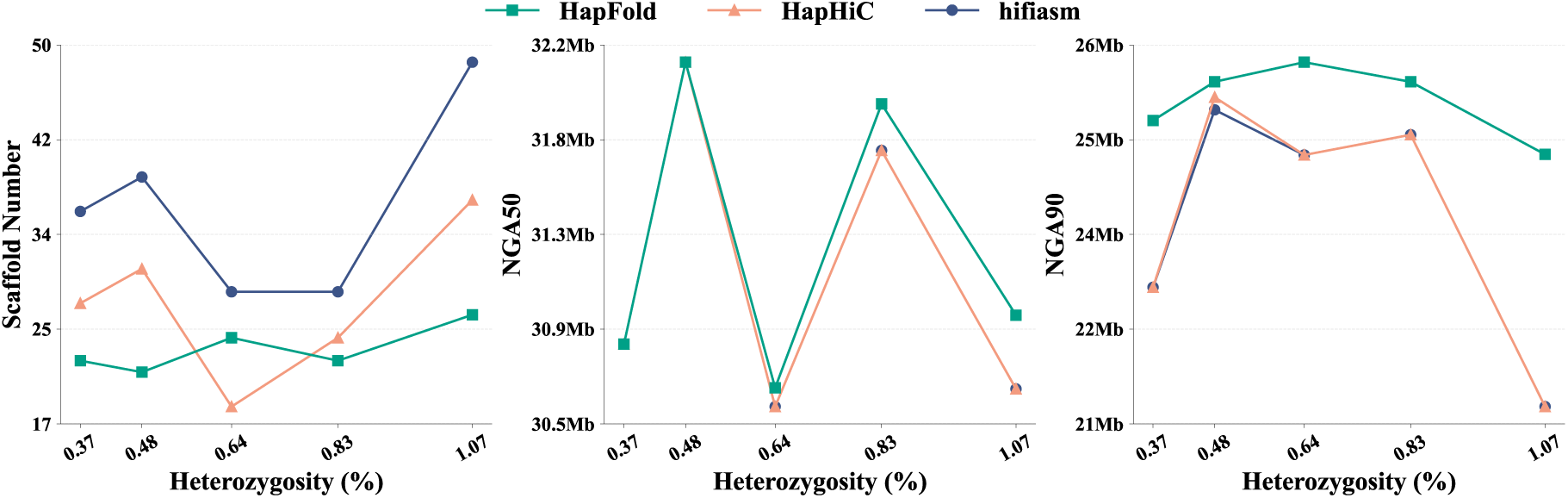
Performance comparison of HapFold, HapHiC, and hifiasm across varying levels of heterozygosity. Different heterozygosity levels were generated by pairwise combinations of distinct rice genomes. The three panels (left to right) show Scaffolds Number, NGA50, and NGA90, respectively. Each curve represents one method, illustrating its performance trend as heterozygosity increases. HiFi reads were simulated using PBSIM2 with an accuracy of 99.87%, and Hi-C paired-end reads were generated using sim3C.

**Extended Data Fig. 5.**
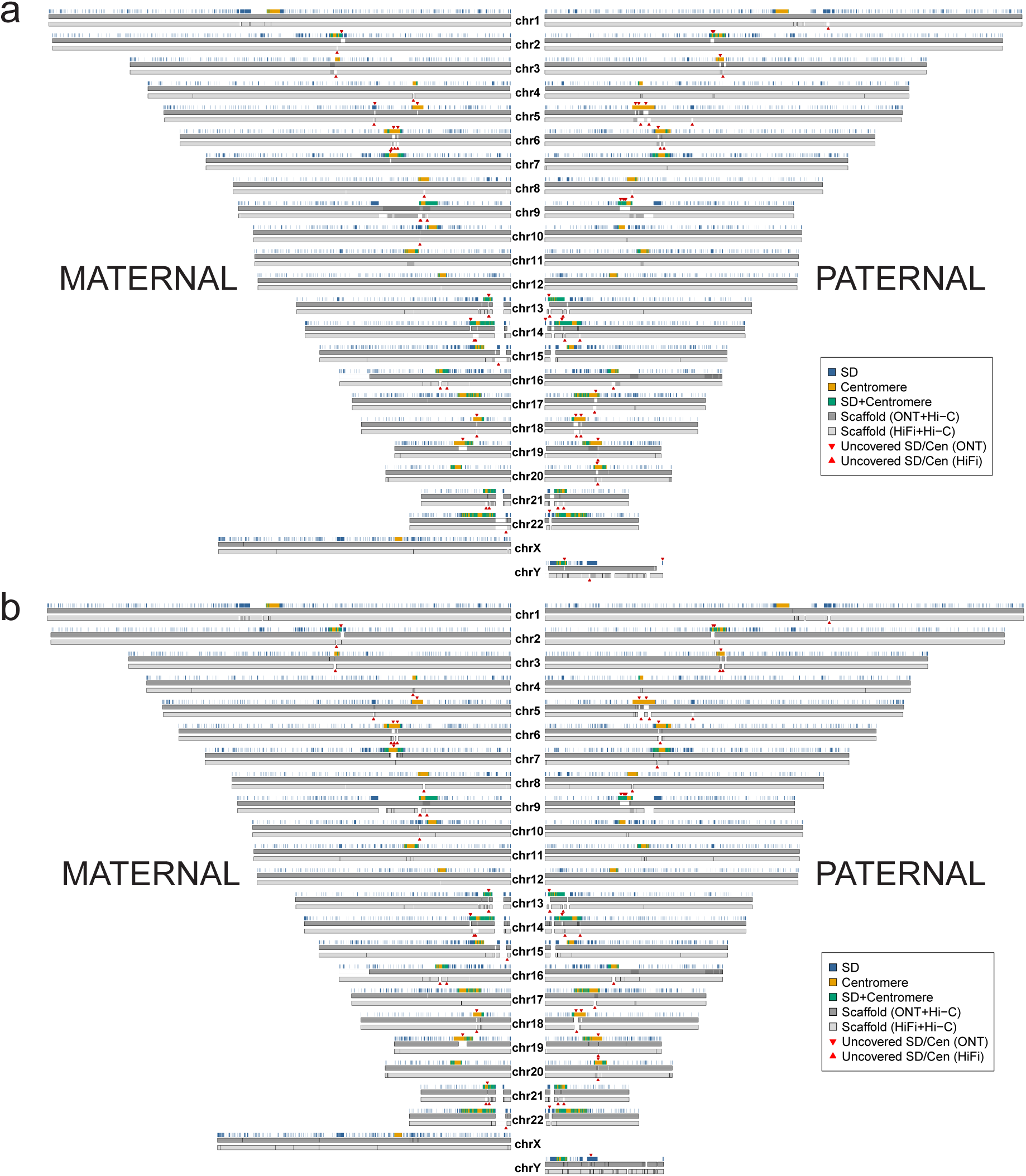
Genome-wide evaluation and gap distribution of HG002 assemblies using PacBio HiFi and standard ONT datasets. All assemblies were accurately assessed against the HG002 Q100 reference genome. Segmental duplication (SD) and centromere (Cen) annotations are shown in blue and yellow, respectively, with overlapping regions highlighted in green. Note that only assembly gaps exceeding 100 kb are defined as uncovered SD/Cen regions. Sites where the assemblies fail to cover these SD/Cen regions are marked with red triangles. a, Genome-wide distribution of the HapFold assemblies and uncovered SD/Cen regions. b, Genome-wide distribution of the hifiasm assemblies and uncovered SD/Cen regions.

**Extended Data Fig. 6.**
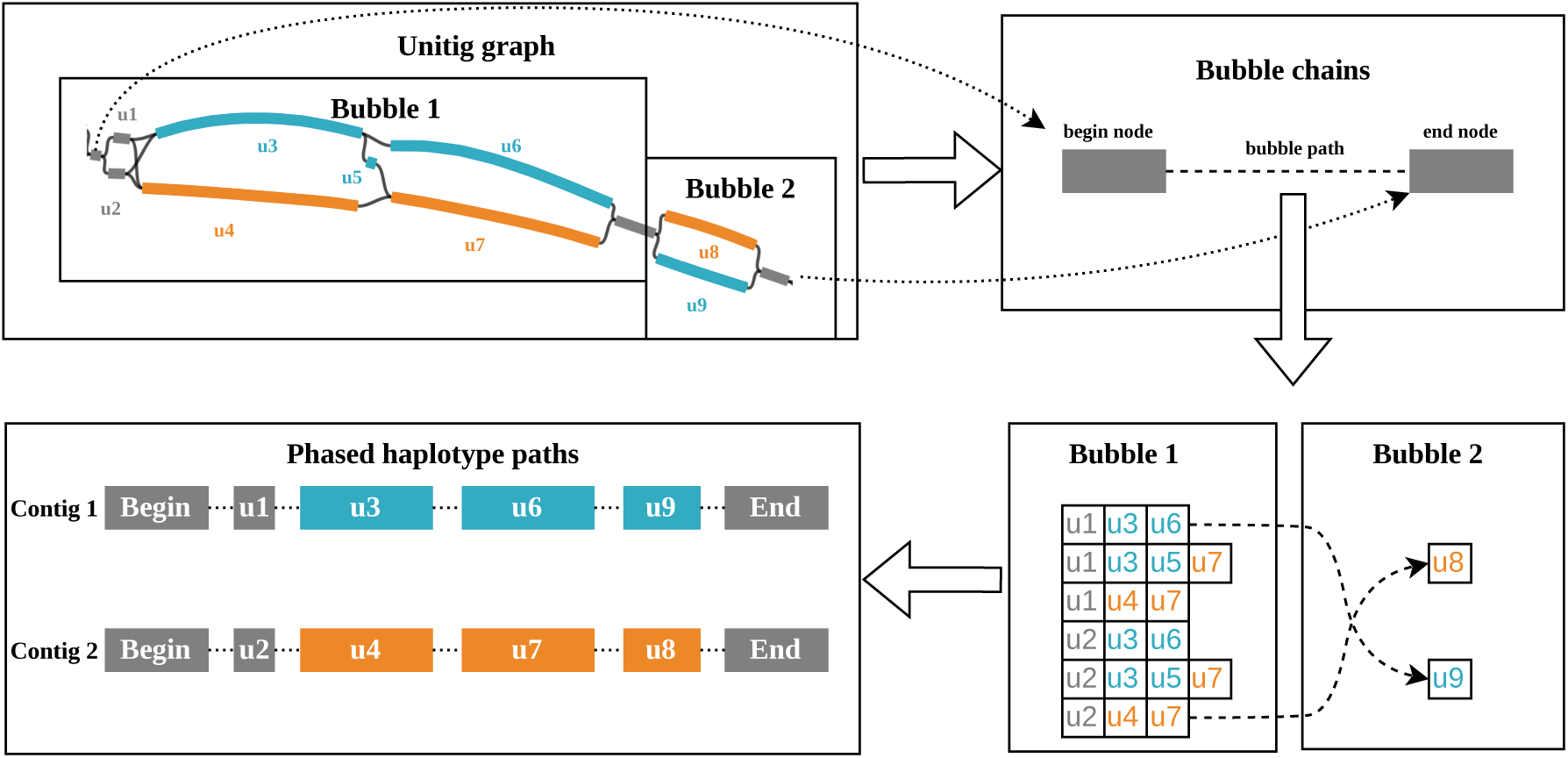
Bubble chains phasing. A local unitig graph contains two bubbles, with cyan and orange paths representing the two parental haplotypes. Graph refinement converts connected bubbles into bubble chains defined by begin and end nodes, while retaining internal bubble paths for phasing. Bubble paths are phased within and across bubbles using Hi-C or Pore-C chromatin contact signals, producing paired phased haplotype paths corresponding to Contig 1 and Contig 2.

**Extended Data Fig. 7.**
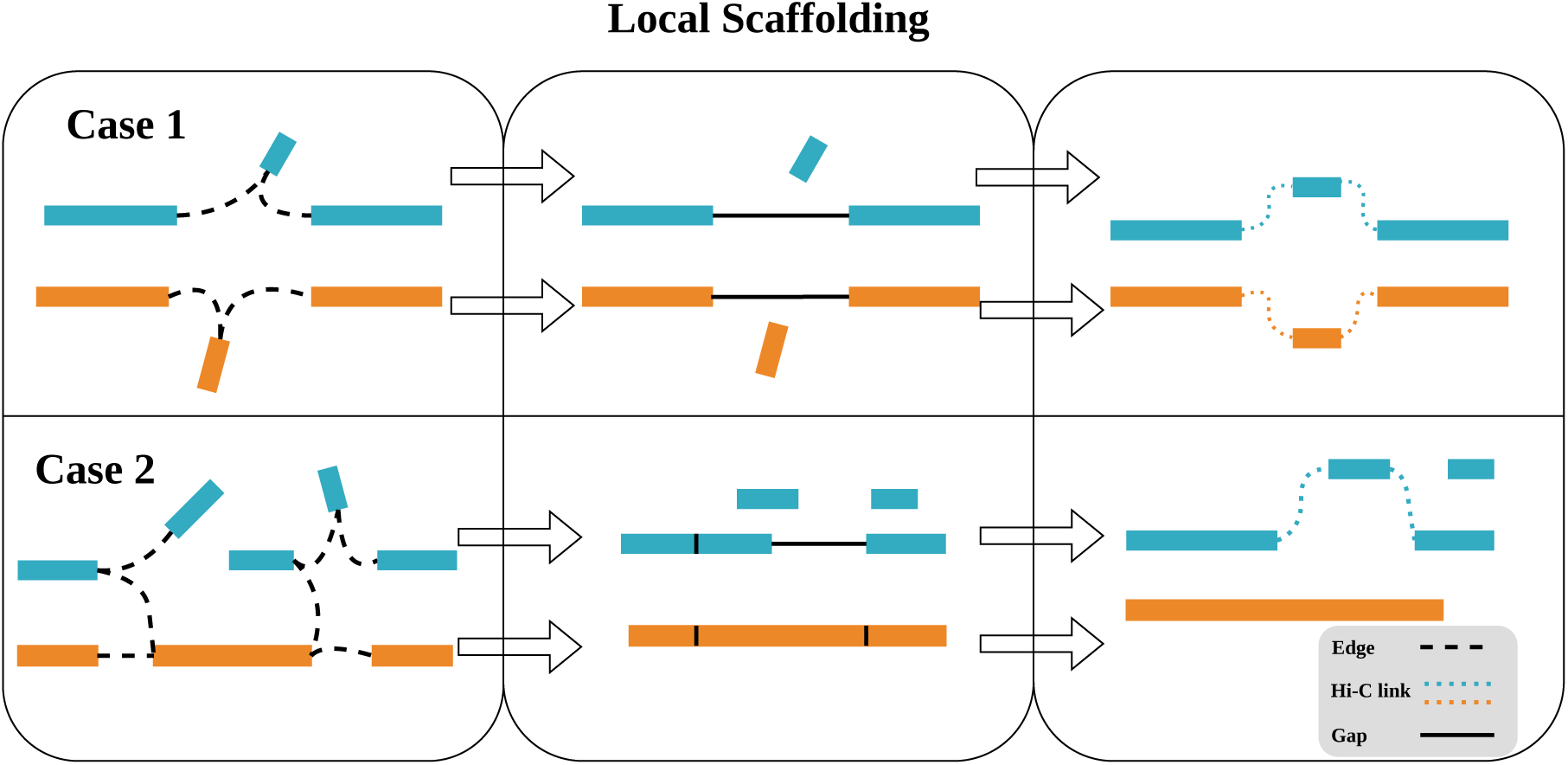
Local scaffolding. Schematic illustration of local scaffolding after haplotype phasing. Cyan and orange segments denote contigs assigned to the two haplotypes. Case 1 and Case 2 illustrate two representative graph configurations that may arise after phasing and the corresponding workflow by which local scaffolding is performed. Solid black lines indicate gap-containing joins between contigs, whereas black dashed lines indicate graph edges supported by overlap. Coloured dashed lines denote Hi-C- or Pore-C-supported linkage signals used to guide haplotype-consistent scaffolding. Vertical black ticks within orange or cyan segments indicate contigs joined without an intervening gap.

**Extended Data Fig. 8.**
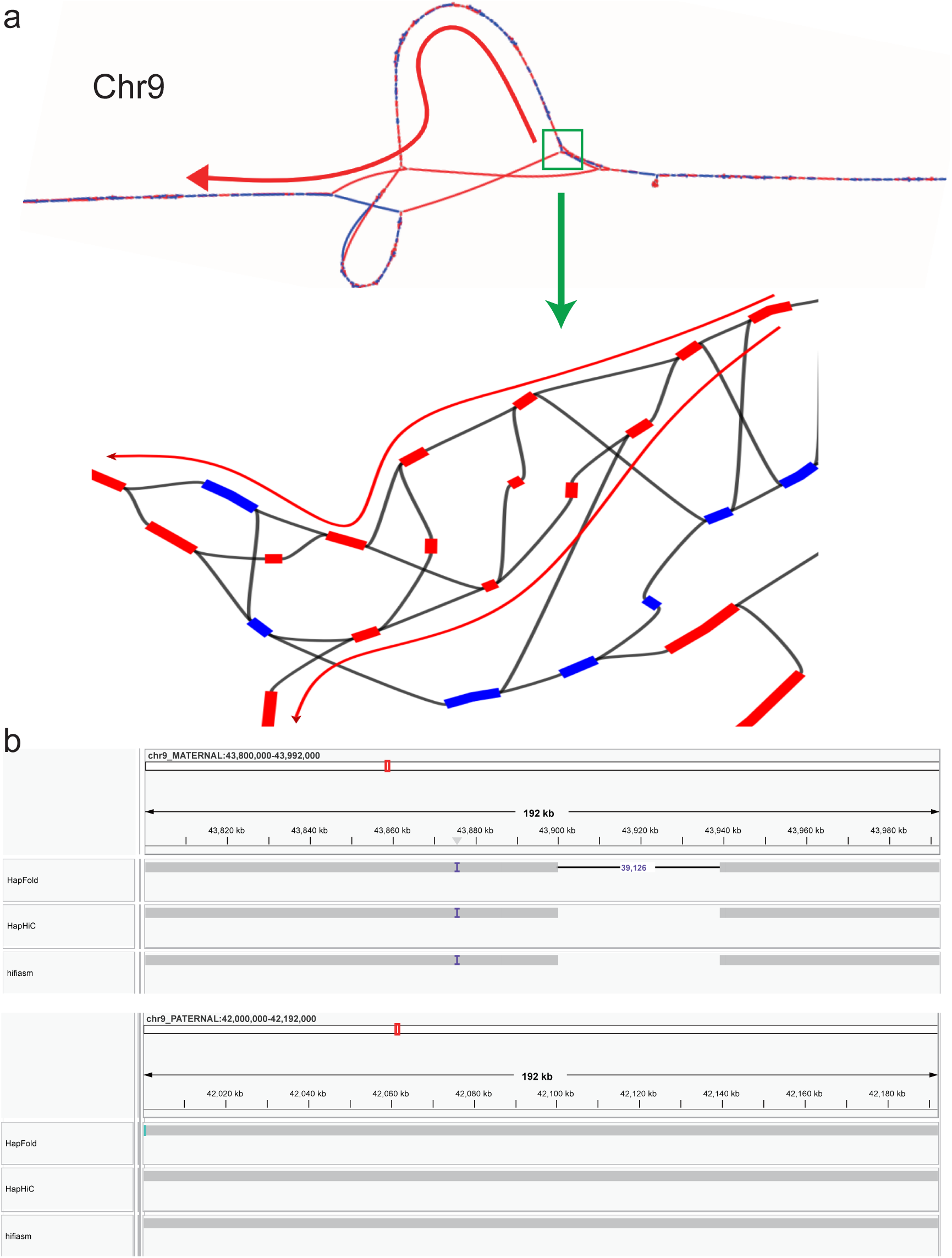
Resolution of complex tangled regions. a, Visualization of the unitig graph connectivity corresponding to a complex region on Chromosome 9 (Chr9 MAT: 43.8–43.9 Mb). Nodes are colored by haplotype origin, with red representing maternal sequences and blue representing paternal sequences. Within this specific region, the maternal haplotype exhibits highly complex branching paths, forming a tangled graph structure. b, Integrative Genomics Viewer (IGV) alignment tracks for the corresponding maternal (top) and paternal (bottom) regions, comparing the assemblies generated by HapFold, HapHiC, and hifiasm. On the maternal track, the structural breaks (blank spaces) in the HapHiC and hifiasm alignments indicate that the assembly is fragmented into two separate sequences at this locus. In contrast, the connected line in the HapFold track demonstrates that it successfully bridged the tangled region, maintaining scaffold contiguity across a defined sequence gap. On the paternal track, the alignments confirm that all three tools completely and accurately resolved the corresponding region without fragmentation.

