## Supplementary Material for "Efficient and accurate near telomere-to-telomere haplotype reconstruction of diploid genomes"

#### Supplementary Tables and Figures

| Category | Metric | HG002 | Chicken1 | Chicken2 | Pig |
|---|---|---|---|---|---|
| **Genome** | Genome size (Gb) | 6.00 | 2.14 | 2.11 | 5.05 |
| | Het Rate (%) | 0.218 | 0.737 | 0.762 | 0.644 |
| | Number of chrs (2n) | 46 | 78 | 78 | 38 |
| **HiFi** | N50 (bp) | 16918 | 18060 | 21262 | 20004 |
| | Total Bases (Gb) | 194.34 | 95.33 | 60.30 | 110.57 |
| | Coverage (×) | 64 | 90 | 58 | 44 |
| **Hi-C** | Total Bases (Gb) | 221.07 | 260.17 | 116.45 | 142.10 |
| | Coverage (×) | 74 | 244 | 110 | 56 |
| **ONT simplex reads** | Reads ≥100 kb (Gb) | 1.47 | - | - | |
| | N50 (bp) | 29395 | - | - | - |
| | Total Bases (Gb) | 153.68 | - | - | - |
| | Coverage (×) | 51 | - | - | - |

Table 1: **Genome characteristics and sequencing data.** The table outlines the fundamental genomic features and raw sequencing metrics for the four diverse biological samples (HG002, Chicken1, Chicken2, and Pig) evaluated in this study. Performance metrics include the read length N50, total sequence yield (Total Bases), and the estimated depth of coverage (×) relative to the estimated genome size. Note that for the Chicken1 sample, Pore-C sequencing data was generated and utilized in place of conventional Hi-C data for chromosomal scaffolding.

| Dataset | HapFold | hifiasm | pstools | HapHiC | YaHS | SALSA2 | 3D-DNA |
|---|---|---|---|---|---|---|---|
| HG002 HiFi+Hi-C | 139.97 | 84.32 | 34.07 | 143.59 | 160.91 | 89.20 | - |
| Chicken1 HiFi+Pore-C | 29.06 | 20.26 | 20.12 | 38.03 | 74.54 | 29.06 | 5.35 |
| Chicken2 HiFi+Hi-C | 29.31 | 14.79 | 17.77 | 38.49 | 83.79 | 20.06 | 3.41 |
| Pig HiFi+Hi-C | 109.21 | 35.99 | 78.79 | 141.21 | 159.74 | 70.64 | 24.17 |
| HG002 ONT+Hi-C | 146.69 | 133.58 | 38.60 | 135.69 | 154.40 | 134.00 | 133.58 |

Table 2: **N50 performance comparison of Hi-C-based scaffolding methods.** Each value represents the diploid N50 metric (in Mb) achieved by a specific scaffolding tool on the corresponding dataset. The N50 is defined as the sequence length of the shortest scaffold at 50% of the total assembly size.

| Dataset | Assembler | Size (GB) | # of Scaffold | N50 (Mb) | NGA50 (Mb) | GF (%) | DR (%) | # of MS | HE (%) | SE (%) |
|---|---|---|---|---|---|---|---|---|---|---|
| HG002(HiFi + 28× Hi-C) | HapFold | 5.97 | 140 | 139.31 | 96.26 | 97.02 | 1.03 | 61 | 3.48 | 0.96 |
| | hifiasm | 5.99 | 627 | 82.13 | 79.48 | 99.48 | 1.01 | 78 | 1.44 | 0.97 |
| | pstools | 9.55 | 850 | 34.11 | 49.97 | 89.408 | 1.76 | 97 | 1.37 | 0.92 |
| | HapHiC | 5.99 | 462 | 145.52 | 87.13 | 99.47 | 1.00 | 83 | 12.76 | 0.98 |
| | YaHS | 5.99 | 228 | 160.78 | 91.58 | 98.71 | 1.01 | 79 | 14.57 | 0.98 |
| | SALSA2 | 5.99 | 535 | 89.43 | 84.31 | 99.47 | 1.01 | 109 | 4.42 | 0.97 |
| HG002(HiFi + 54× Hi-C) | HapFold | 5.97 | 137 | 147.17 | 93.89 | 98.21 | 1.01 | 68 | 3.41 | 0.96 |
| | hifiasm | 5.99 | 634 | 80.54 | 73.13 | 99.30 | 1.01 | 77 | 1.443 | 0.98 |
| | pstools | 9.55 | 850 | 34.11 | 49.96 | 89.61 | 1.76 | 97 | 1.37 | 0.92 |
| | HapHiC | 5.99 | 463 | 146.88 | 89.42 | 99.30 | 1.01 | 76 | 4.08 | 0.98 |
| | YaHS | 5.99 | 252 | 160.21 | 89.78 | 98.03 | 1.02 | 85 | 14.56 | 0.98 |
| | SALSA2 | 5.99 | 549 | 89.43 | 82.13 | 99.29 | 1.01 | 107 | 3.57 | 0.98 |
| HG002(HiFi + 76× Hi-C) | HapFold | 5.97 | 122 | 135.52 | 99.12 | 98.81 | 1.01 | 40 | 4.26 | 0.97 |
| | hifiasm | 5.99 | 631 | 84.32 | 64.84 | 99.39 | 1.01 | 114 | 1.54 | 0.97 |
| | pstools | 8.97 | 826 | 34.07 | 49.46 | 88.23 | 1.68 | 136 | 1.35 | 0.91 |
| | HapHiC | 5.99 | 426 | 143.59 | 91.86 | 99.52 | 1.00 | 67 | 6.88 | 0.97 |
| | YaHS | 5.99 | 269 | 160.91 | 74.20 | 99.49 | 1.00 | 99 | 14.90 | 0.97 |
| | SALSA2 | 5.99 | 540 | 89.20 | 64.84 | 99.39 | 1.01 | 147 | 3.94 | 0.97 |
| HG002(HiFi + 100× Hi-C) | HapFold | 5.97 | 136 | 139.94 | 94.81 | 98.41 | 1.01 | 61 | 2.65 | 0.95 |
| | hifiasm | 5.99 | 651 | 82.08 | 74.21 | 99.07 | 1.01 | 68 | 1.39 | 0.97 |
| | pstools | 9.55 | 846 | 34.92 | 49.96 | 89.18 | 1.77 | 94 | 1.37 | 0.92 |
| | HapHiC | 5.99 | 448 | 139.99 | 91.06 | 99.59 | 1.00 | 64 | 6.00 | 0.97 |
| | YaHS | 5.99 | 288 | 146.90 | 91.69 | 99.50 | 1.00 | 82 | 14.46 | 0.97 |
| | SALSA2 | 5.99 | 568 | 87.31 | 86.97 | 99.04 | 1.01 | 94 | 3.94 | 0.97 |

Table 3: **Performance Evaluation of Hifiasm Assembly and Scaffolding Tools across Varying Hi-C Sequencing Depths for HG002.** All metrics are reported for the diploid assembly (sum of both haplotypes). Size: total assembled diploid genome size. # of scaffold: the total number of assembled sequences. N50: the length of the shortest scaffold at 50% of the total assembly size. NGA50: N50 calculated using reference-aligned blocks, providing a robust measure of structural continuity and accuracy. GF (genome fraction): percentage of the reference genome successfully covered by the assembly. DR (duplication ratio): ratio of total aligned bases in the assembly to the total covered bases in the reference. # of MS (misassembled scaffolds): number of scaffolds containing structural assembly errors (relocations, inversions, or translocations). N50, NGA50, genome fraction, duplication ratio, and number of misassembled scaffolds were comprehensively evaluated using the tool QUAST[1] based on alignments to corresponding reference genomes. HE (hamming error): proportion of incorrectly phased alleles within phase blocks. SE (switch error): rate of phase switches between adjacent heterozygous loci. Both phasing metrics were evaluated using the $k$-mer-based tool yak.

| Dataset | Assembler | QV |
|---|---|---|
| HG002(HiFi + Hi-C) | HapFold | 39.25/44.40 |
| | hifiasm | 39.24/44.35 |
| | HapHiC | 39.22/44.31 |
| HG002(ONT + Hi-C) | HapFold | 38.66/42.61 |
| | hifiasm | 38.65/42.61 |
| | HapHiC | 38.65/42.61 |

Table 4: **Assembly Quality Value (QV) Comparison of Different Assemblers on HG002 Datasets.** The table presents the QV scores for three assemblers—HapFold, hifiasm, and HapHiC—using two distinct sequencing combinations: HG002 (HiFi + Hi-C) and HG002 (ONT + Hi-C). These QV scores, which reflect per-base assembly accuracy, were evaluated using the k-mer-based tool yak). Both values are provided by yak and presented in the format of *raw_quality_value / adjusted_quality_value*.

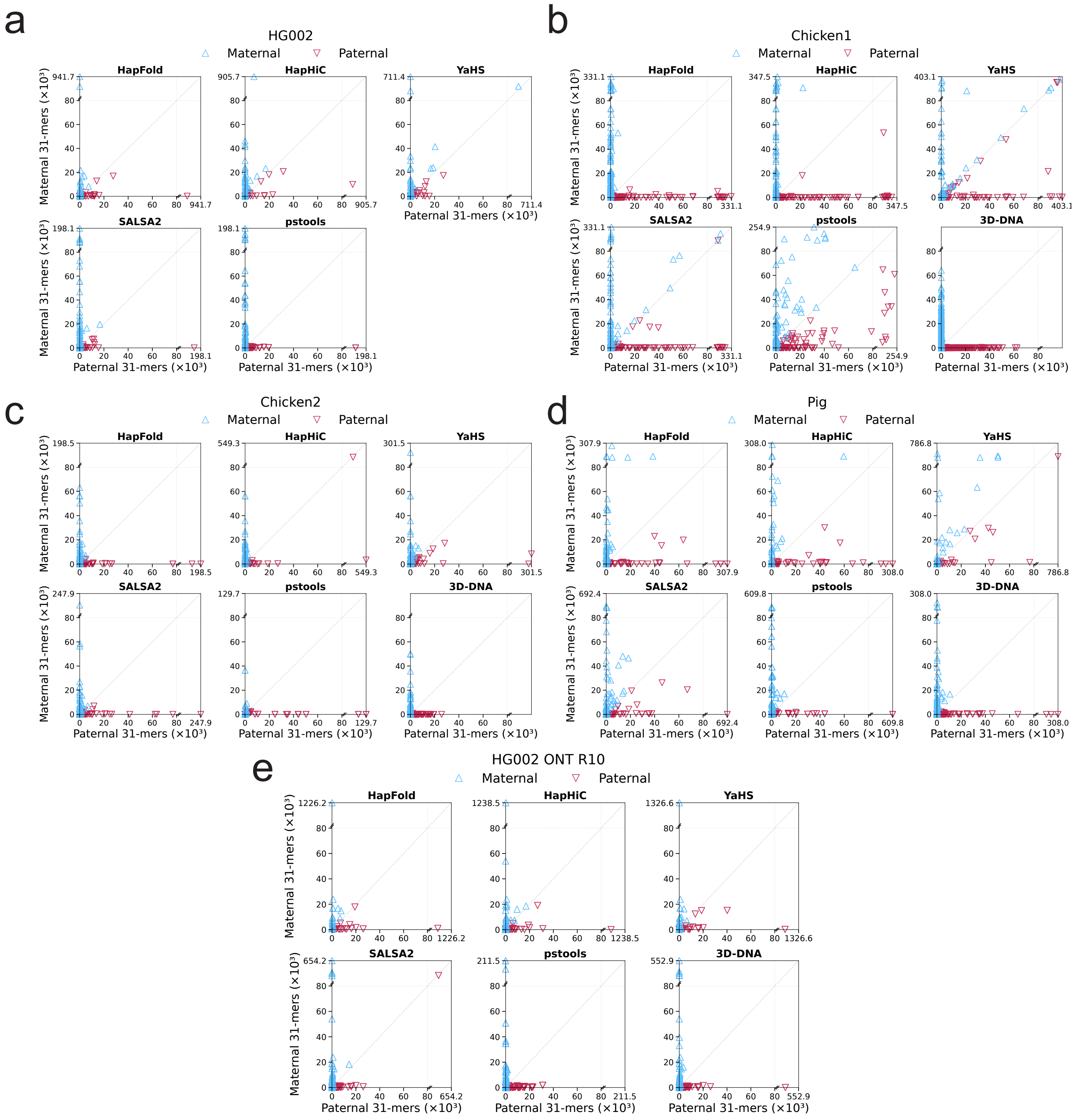


Figure 1: **Evaluation of haplotig assembly accuracy across different assemblers and datasets.** The figure displays k-mer completeness and switch error analysis for various genome assemblies (HapFold, HapHiC, YaHS, SALSA2, ptools, and 3D-DNA). Assemblies were evaluated using paternal and maternal 31-mer k-mer counts across four distinct datasets: the human HG002 genome, two chicken genomes (Chicken1 and Chicken2), and a pig genome. Additionally, the bottom panel specifically evaluates the HG002 assembly using Oxford Nanopore (ONT) R10 sequencing data. Each scatter plot represents the distribution of assembled contigs based on their inheritance, where triangles pointing upward ($\triangle$) and downward ($\bigtriangledown$) denote maternal and paternal haplotigs, respectively. Axes are broken at a threshold of 80 $\times 10^3$ k-mers to visualize both high-coverage contigs and low-abundance assembly artifacts. The dashed diagonal line indicates the expected 1:1 ratio for homozygous/balanced regions.

### Software Commands

This section summarizes only the core computational commands used in the workflows.

#### 1 Data Simulation

##### 1.1 PacBio HiFi read simulation using pbsim2

pbsim2 (`https://github.com/yukiteruono/pbsim2`) was used to simulate PacBio HiFi reads from the reference genome. The simulator allows generating highly accurate long reads with configurable length distribution and error profiles. The command used in this study is as follows:

```
pbsim2 --depth 20 --sample-fastq <sample/sample.fastq> <sample/sample.fasta>
```

##### 1.2 Hi-C read simulation using sim3C

sim3C (`https://github.com/ivargr/sim3C`) was used to simulate Hi-C contact reads based on the reference genome. This tool models chromatin conformation capture experiments and generates paired-end reads reflecting spatial proximity in the genome. The command used in this study is as follows:

```
sim3C --dist uniform -n <pairs> -l 150 -e <enzyme> -m hic <myref.fasta> 1.fq 2.fq
```

#### 2 Genome assembly

##### 2.1 hifiasm

hifiasm (`https://github.com/chhylp123/hifiasm`, commit: ec9a8b2) is a de novo assembler for PacBio HiFi reads. It supports multiple assembly modes depending on available sequencing data. The following commands demonstrate the three main modes used in this study:

**(1) Haplotype-resolved assembly using PacBio HiFi long reads and Hi-C contact data**

```
hifiasm -o <asm> -t 32 --h1 <1.fq> --h2 <2.fq> <HiFi.fq>
```

**(2) Haplotype-resolved assembly combining PacBio HiFi, Oxford Nanopore UltraLong reads, and Hi-C data**

```
hifiasm -o <asm> -t 32 --h1 <1.fq> --h2 <2.fq> --ul <ONT.fq> <HiFi.fq>
```

**(3) Haplotype-resolved assembly using Oxford Nanopore R10 long reads and Hi-C data without PacBio sequences**

```
hifiasm -o <asm> -t 32 --ont --dual-scaf --telo-m CCCTAA --h1 <1.fq> --h2 <2.fq> <ONT_R10.fq>
```

#### 3 Quality control, mapping, and filtering of Hi-C reads

##### 3.1 General purpose Hi-C read mapping and filtering

The following two command blocks are from the HapHiC(`https://github.com/zengxiaofei/HapHiC`, commit: df0f5ac) preprocessing workflow (HapHiC toolkit and its `utils` scripts).

**(1) First, BWA-MEM (0.7.18-r1243-dirty) with the "-5SP" parameter was used to align Hi-C reads, samblaster (version 0.1.26) was used to mark PCR duplicates, and SAMtools (version 1.21) with flag "3340" was used to remove secondary and supplementary alignments.**

```
bwa index <asm.fa>
bwa mem -5SP -t 32 <asm.fa> <1.fq> <2.fq> | samblaster | samtools view -@ 16 -S -h -b -F 3340 -o <HiC.bam>
```

**(2) Then, HapHiC-provided `filter_bam.py` (from the `utils` directory of the HapHiC GitHub repository) was used to remove alignments with mapping quality zero ($MAPQ = 0$) and edit distance greater than 2 ($NM \geq 3$).**

```
python3 filter_bam.py <HiC.bam> 1 --nm 3 --threads 32 | samtools view -b -@ 16  -o <HiC.filtered.bam>
```

##### 3.2 Hi-C read mapping and filtering for 3D-DNA scaffolding

We used the Juicer pipeline (`https://github.com/aidenlab/juicer`, commit: f866383) to align and filter Hi-C reads for 3D-DNA as follows:

###### (1) prepare scripts

```
ln -s /path/to/juicer1.6/CPU scripts
```

###### (2) prepare reads

```
mkdir fastq && cd fastq
ln -s <1.fq> reads_R1.fastq.gz && ln -s <2.fq> reads_R2.fastq.gz && cd ..
```

###### (3) prepare genome

```
mkdir references && cd references
ln -s <asm.fa> genome.fa && bwa index genome.fa && cd ..
```

###### (4) prepare restriction sites

```
python2 /path/to/juicer1.6/misc/generate_site_positions.py DpnII genome ../references/genome.fa
awk 'BEGIN{OFS=``\t''}{print $1, $NF}' genome_DpnII.txt > genome.chrom.sizes
```

###### (5) run Juicer pipeline

```
/path/to/juicer1.6/CPU/juicer.sh -g genome -z references/genome.fa -p restriction_sites/genome.chrom.sizes -y
restriction_sites/genome_DpnII.txt -D `pwd` -s DpnII -t 32
```

#### 4 Hi-C-based scaffolding

##### 4.1 HapFold

HapFold (`https://github.com/LuoGroup2023/HapFold`, commit: 83997c8)

```
awk '/^S/{print ``>'' $2; print $3}' <asm.hic.p_utg.gfa> > asm.hic.p_utg.gfa.fa
HapFold mapping -t 32 asm.hic.p_utg.gfa.fa <1.fq> <2.fq> -o <gfa_hic.out>
```

```
HapFold scaffolding -t 32 -n <nchrs> -u <utg_ctg_mappings.csv> -i true <gfa_hic.out> <asm.hic.p_utg.gfa> <out>
-1 <asm.hic.hap1.p_ctg.gfa> -2 <asm.hic.hap2.p_ctg.gfa>
```

##### 4.2 HapHiC

HapHiC (`https://github.com/zengxiaofei/HapHiC`, commit: df0f5ac) was typically executed using default parameters without assembly correction:

###### (1) Default run: HapHiC was first executed with default parameters for standard clustering and scaffolding.

```
haphic pipeline <asm.fa> <HiC.filtered.bam> <nchrs>
```

###### (2) If clustering was unsatisfactory, HapHiC was re-run with adjusted inflation-related parameters to improve clustering behavior.

```
haphic pipeline <asm.fa> <HiC.filtered.bam> <nchrs> --max_inflation 10 --inflation_step 0.2
```

##### 4.3 YaHS

YaHS (`https://github.com/c-zhou/yahs`, commit: 9b2cc15) was typically executed without assembly correction and memory check:

```
samtools faidx <asm.fa>
yahs <asm.fa> <HiC.bam> -q 1 --no-contig-ec --no-mem-check
```

##### 4.4 SALSA2

SALSA2 (`https://github.com/marbl/SALSA`, commit: 1b76bf6) was executed without an assembly graph.

**(1) Assembly correction was typically disabled. Given that the BAM file "HiC.bam" is already sorted by name, the sorting step was skipped:**

```
samtools faidx <asm.fa>
bedtools bamtobed -i <HiC.bam> > alignment.bed
python2 run_pipeline.py -a <asm.fa> -l <asm.fa.fai> -b alignment.bed -e <RE> -m no -o out
```

**(2) For scaffolding assemblies with chimeric contigs, assembly correction was enabled:**

```
python2 run_pipeline.py -a <asm.fa> -l <asm.fa.fai> -b alignment.bed -e <RE> -m yes -o output
```

##### 4.5 pstools

pstools (`https://github.com/shilpagarg/pstools`, commit: 90c1986) was typically executed using default parameters without assembly correction:

```
awk '/^S/{print ``>'' $2; print $3}' <asm.hic.p_utg.gfa> > asm.hic.p_utg.gfa.fa
pstools hic_mapping -t32 -o map.out <asm.hic.p_utg.gfa.fa> <1.fq> <2.fq>
pstools resolve_haplotypes -t32 -i true map.out <asm.hic.p_utg.gfa> out
```

##### 4.6 3D-DNA

3D-DNA (`https://github.com/aidenlab/3d-dna`, commit: cb63403) was typically executed using default parameters without assembly correction:

```
run-asm-pipeline.sh -r 0 <asm.fa> <merged_nodups.txt>
```
